# Clinically driven knowledge distillation for sparsifying high-dimensional multi-omics survival models

**DOI:** 10.1101/2022.02.07.479388

**Authors:** David Wissel, Daniel Rowson, Valentina Boeva

**Affiliations:** Department of Computer Science, ETH Zurich, Zurich, Switzerland; Swiss Institute for Bioinformatics (SIB), Zurich, Switzerland; Institut Cochin, Inserm U1016, CNRS UMR 8104, Paris Descartes University UMR-S1016, 75014 Paris, France

## Abstract

Recently, various methods have been proposed to integrate different heterogeneous high-dimensional genomic data sources to predict cancer survival, often in addition to widely available and highly predictive clinical data. Although clinical applications of survival models have high sparsity requirements, most state-of-the-art models do not naturally exhibit this sparsity, as they are based on random forests or deep learning. We propose to use *𝓁*_1_-penalized linear student models within a knowledge distillation framework to sparsify underlying multi-omics black-box teachers. We show that by excluding clinical variables from our *𝓁*_1_ penalty, we can effectively guide the knowledge distillation, reaching virtually identical discriminative performance to the teachers while using on average 140 features or less across the 17 cancer datasets from The Cancer Genome Atlas (TCGA) considered in our study.

## 1. Introduction

Accurately estimating patient-specific risk is useful in clinical settings for cancer patient stratification and thus therapy choice among other things.^1^ In addition, the accurate identification of risk factors for a particular cancer can drive an improved understanding of upstream causes, and, in turn, treatments. Thus, it is crucial to not only improve the discriminative power of survival models but also to identify the features driving their predictions. While various models have been developed specifically to predict cancer survival from high-dimensional multi-omics data, the best-performing models have not exhibited sparsity (Hornung & Wright, 2019b; Vale-Silva & Rohr, 2021; Cheerla & Gevaert, 2019).

Thus, there is a strong need for methods that can maintain the high performance of deep learning or adapted random forest methods, yet provide relatively strong sparsity in input feature space. We propose to use knowledge distillation (Hinton et al., 2015) using *𝓁*_1_-penalized linear students for sparsifying well-performing black-box high-dimensional multi-omics survival models. Our key contributions were as follows:

- We showed that knowledge distillation could be made effective in survival settings by using the predicted risk measures of teachers as response-based knowledge and excluding clinical variables from our *𝓁*_1_ penalty. Specifically, we explored relevant applications of our proposed distillation approach on The Cancer Genome Atlas (TCGA), revealing that clinically guided knowledge distillations allowed for the sparsification of black-box high-dimensional multi-omics survival models at virtually no loss in discriminative performance relative to the teacher models.
- Further, we showed that the good performance of our students could not be explained merely through feature selection, suggesting the presence of additional effects inherent in the distillation process with survival data.

## 2. Related work

### Black-box multi-omics cancer survival methods

In recent years, there has been a considerable amount of work on survival methods that aim to integrate multiple genomic data sources, so-called multi-omics data. Hornung & Wright (2019b) investigated multiple random forest adaptations, all of which changed the split-point selection of the standard random survival forest model (Ishwaran et al., 2008). The best performing new method introduced by the authors, BlockForest, was shown to significantly outperform a standard random survival forest in terms of Harrell’s concordance across 20 TCGA datasets when integrating multiple omics modalities, in addition to clinical data. Herrmann et al. (2021) further validated BlockForest, showing that it performed the best out of various considered statistical multi-omics models across 18 TCGA datasets and was one of the only models that (marginally) outperformed the baseline of a clinical-only Cox PH model when set to integrate clinical data in addition to multiple omics modalities.

There has also been strong interest in leveraging deep learning for multi-omics integration in cancer survival models. Cheerla & Gevaert (2019) showed that training modality-specific networks that were integrated using mean-pooling could achieve excellent discriminative performance when using clinical data in addition to multiple genomic modalities and whole slide images. Further, Cheerla & Gevaert (2019) revealed that training neural networks using multiple cancers jointly (*i*.*e*., pan-cancer) could produce a distinct benefit in discriminative performance for some but not all cancers. Recently, Vale-Silva & Rohr (2021) showed that adopting a non-proportional hazards loss could further boost the performance of a pan-cancer neural model. The authors showed that their model, which was based on integrating modality-specific representations using max-pooling, could outperform both standard methods such as the Cox PH model and a random survival forest, as well as modern neural models such as *DeepSurv* (Katzman et al., 2018) and *DeepHit* (Lee et al., 2018) when limited to one input modality and trained on all TCGA cancers jointly. In addition, the authors achieved further increases in performance when integrating genomic data in addition to clinical data, with their model performing best when set to integrate clinical data and gene expression.

Wissel et al. (2022) investigated the integration of high numbers of omics modalities, in addition to clinical data, through neural models. Integrating many modalities can be challenging for many models, due to the low predictive power and high noise of some molecular modalities. The authors showed that, for the right choice of modalities (only clinical data and gene expression), the chosen integration technique to unify modality-specific representations (*e*.*g*., mean-pooling) did not matter, with all investigated techniques performing similarly. Meanwhile, when set to integrate all available modalities, an autoencoder outperformed other integration methods for fusing modality-specific representations, seemingly due to its ability to focus on more informative modalities. In addition, the authors showed that using a hierarchical autoencoder for multi-omics integration performed comparably to a variant of BlockForest, RandomBlock favoring clincal variables.

### Sparse linear multi-omics cancer survival methods

Boulesteix et al. (2017) proposed a modified *𝓁*_1_ penalized Cox PH model which used separate regularization hyperparameters λ_1_, …, λ_*m*_ per modality which allowed the model to focus more strongly on some modalities. The authors showed that their newly proposed model, termed IPF-LASSO, outperformed the standard Cox PH Lasso in simulations and two cancer data sets from TCGA and the Gene Expression Omnibus (GEO), each containing clinical data and one or more molecular modalities, in terms of the integrated Brier score. Klau et al. (2018) proposed a sequential approach to integrate multiple modalities in linear sparse cancer survival models. Their proposed method, *prioritylasso*, fitted each modality using an *𝓁*_1_-regularized Cox PH model sequentially and at each step used the predictions from the previous round as an unregularized offset. The authors showed that *prioritylasso* performed as well or better than a Cox PH Lasso model on a cancer survival dataset obtained from GEO that contained both clinical data and molecular modalities. In addition, *prioritylasso* provided clinicians the ability to influence the model by assigning their preferred modality ordering. The model was in turn more likely to include variables belonging to a block that had been assigned a higher priority.

### Knowledge distillation

Knowledge distillation was coined by Hinton et al. (2015), who showed that, in a classification setting, the performance of an ensemble of teacher models could be matched by a single student model that was trained using a combination of the original labels and the logits predicted by the teachers. Since its introduction, knowledge distillation has found countless applications to, among others, language models (Sun et al., 2019), domain adaptation (Nguyen-Meidine et al., 2021; Zhang et al., 2021) and various other problems. Nevertheless, despite recent advances (Stanton et al., 2021; Phuong & Lampert, 2019; Menon et al., 2021; Yuan et al., 2020), there is still large amounts of ambiguity as to why knowledge distillation works, even in the classification setting. In non-classification settings, knowledge distillation is exceedingly rare (Saputra et al., 2019). To the best of our knowledge, no previous work has explored knowledge distillation within a survival analysis setting.

## 3. Clinically guided knowledge distillation

We propose to use knowledge distillation (KD) for sparsifying black-box multi-omics survival models. We considered the simplified setting of an *𝓁*_1_-penalized linear student model. In the survival setting, in contrast to classification, there is no temperature hyperparameter. Further, to make distillation applicable to arbitrary risk measures, we set *α* = 0 in the common KD setting (Equation (1)). Thus, we discarded survival time and censoring information and instead trained our students only on the knowledge provided by their teachers.

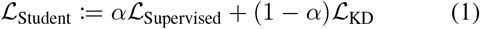

We used a teacher-specific risk measure as the response-based knowledge to be transferred to our students. Given a predicted teacher risk measure 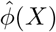, our students were fitted as:

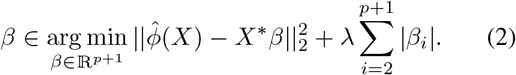

Here, λ was a hyperparameter governing the strength of the *𝓁* _1_ penalty, X ∈ ℝ^*n*×*p*^ was our design matrix, and 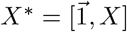 was the design matrix with an intercept vector.

Furthermore, we considered students which did not penalize clinical variables. The idea of not penalizing clinical variables was based on the findings of several studies that in some cancers, molecular variables offered limited added predictive value besides clinical information (Herrmann et al., 2021). In addition, treating clinical variables preferentially (*e*.*g*., through excluding from the *𝓁*_1_ penalty) has shown promise in both linear cancer survival models (Klau et al., 2018; Herrmann et al., 2021) and random forests adapted to multi-omics cancer survival data (Hornung & Wright, 2019b):

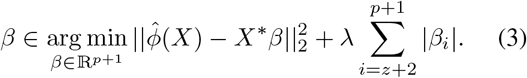

Here, the first z columns of our design matrix X corresponded to clinical variables. We refer to the latter version of considered knowledge distillation as *clinically guided* students or *clinically guided* distillation. We now move on to discussing our experimental setup, through which we validated the effectiveness of our proposed distillation approach.

## 4. Experiments

### 4.1. Datasets

We considered 17 out of the 33 available cancer datasets from TCGA, following the dataset selection of Herrmann et al. (2021). We excluded datasets that contained less than 100 samples *after* preprocessing, datasets that did not have a minimum event ratio of 5%, and datasets that did not have all required molecular modalities. We also excluded Acute Myeloid Leukemia (LAML) relative to the work of Herrmann et al. (2021) since it contained only 35 samples after preprocessing in their study.

We used the same clinical variables as Herrmann et al. (2021) except that we excluded clinical variables that were missing for more than five patients on a particular dataset.^2^

We excluded any patients that were still missing any of the thus chosen clinical variables. Categorical clinical variables were one-hot encoded.

We used all molecular information broadly available on TCGA, except whole slides images. We thus used, in addition to clinical data, gene expression (GEX), mutation, DNA methylation, copy number variation (CNV), reverse phase protein array (RPPA), and miRNA. We excluded patients that did not have all required genomic modalities (that is, patients that were, for example, completely missing the RPPA modality). Molecular features that were missing for more than one patient were excluded to conserve as many patients as possible. We used log-transformed values of miRNA and gene expression. All other molecular input data were used without further preprocessing. We provide a link to our complete datasets within our code repository.

### 4.2. Teacher models

For the considered teacher models, we restricted ourselves to methods that were originally designed for individual cancer datasets, that is, we excluded methods fit on pan-cancer data, both since clinical data are often cancer-specific and since this would have made us deviate from our main research question.

We considered BlockForest (Hornung & Wright, 2019b) (BF), a random forest method adapted to multi-omics data. In addition, we included RandomBlock, a variant of BlockForest, which has been shown to outperform BlockForest if clinical variables were *favored* (mandatorily considered at every split point selection within each tree) (Hornung & Wright, 2019b; Wissel et al., 2022). For RandomBlock, we favored clinical features and will thus refer to it as RandomBlock favoring (RBF). Since both BF and RBF were based on random survival forests, we used the *ensemble mor*tality as its predicted risk measure that was used to transfer knowledge to our students (Ishwaran et al., 2008).

We also included a neural model, HierarchicalSAE (HSAE), based on hierarchical autoencoders, that was shown to perform comparably to RandomBlock favoring, in a recent study (Wissel et al., 2022). Since HSAE optimized a Cox PH loss, we used its patient-specific estimated log-risk to transfer knowledge to our students.

### 4.3. Reference methods

For reference methods, we included a Lasso regularized Cox PH model, for which we did not penalize clinical variables (Lasso favoring). Thus, our first reference method was the same model as our clinically guided students (Equation (3)), except that it was fit on the survival targets as opposed to the teacher knowledge. Second, we considered *prioritylasso*, a sequential Lasso-based Cox PH model (Klau et al., 2018), which has been shown to perform well on multi-omics data when given a good pre-specified priority order (Wissel et al., 2022).

We initially included Coxboost favoring (Binder, 2021), as it was shown to perform well in the study of Herrmann et al. (2021). We found, however, that even when dummy-encoding (as opposed to one-hot encoding) clinical categorical variables, we consistently encountered strong convergence issues for the clinical coefficient estimates, which led to the exclusion of clinical features by the model in most cases. Thus, we found Coxboost favoring to under-perform overall relative to a clinical-only Cox PH model (data not shown). Since we were interested in benchmarking sparse models which could (in specific settings) outperform the clinical-only model, we did not include our results for Coxboost favoring in the main text.

### 4.4. Performance metrics

For measuring student fidelity, that is, the degree to which students were able to match their teachers we considered Spearman’s ρ, the standard rank correlation coefficient (Spearman, 1904). We calculated Spearman’s ρ on both the train and the test set, each time calculating the rank correlation between the teacher predictions and the student predictions.

As not all students could produce predictions of the survival function, we restricted ourselves to Harrell’s concordance (Harrell et al., 1982) (Equation (4)) when evaluating teacher and student generalization performance, the *de facto* standard measure for evaluating the discriminative performance of survival models.^3^ We let 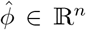 be a vector of estimated, patient-specific risk measures, and let *δ* ∈ 𝔹^*n*^ be the censoring indicator, that is B_*i*_ = 1 iff patient i experienced the event during the study. Lastly, we let 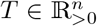 be the set of observed times, that is, for a specific patient *i*, we let *T*_*i*_ ≡ min(*C*_*i*_, *U*_*i*_), where *C*_*i*_ denotes the time at which the patient was censored and *U*_*i*_ denotes the time at which the patient experienced the event.

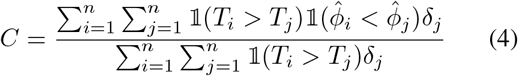

For sparsity, we considered the degree of sparseness of each model, that is, the number of variables that each model used for making predictions.

### 4.5. Validation

For validation, on each dataset, we used five-fold cross-validation, stratified for the censoring indicator, twice repeated, yielding 10 test splits per dataset.

Although pooling performance measures across datasets has been described as questionable (Demšar, 2006), depending on the setting, pooling test concordances across cancers on TCGA is somewhat standard, for both models training on multiple cancers and models trained only on individual cancers (Herrmann et al., 2021; Vale-Silva & Rohr, 2021; Cheerla & Gevaert, 2019; Kim et al., 2020). Thus, for convenience of exposition, we will, at times also pool performance metrics across datasets.

When pooling test concordances across datasets, we included Wilcoxon ranked sign significance tests that have been recommended to compare models across datasets (Demšar, 2006). For the Wilcoxon ranked sign tests, to summarise the performance of each model on each dataset, we calculated the mean concordance across test splits on each dataset and used these means per dataset as the performances to be tested for differences, following the approach of (Hornung & Wright, 2019b). We emphasize that a higher pooled mean concordance does not necessarily correspond to a more significant performance difference as measured by the p-value of the respective Wilcoxon ranked sign test.

We did not test for significant performance differences on individual cancer datasets, as tests adjusting for the repeated cross-validation that we used, such as the corrected repeated k-fold cv test (Bouckaert & Frank, 2004), would have been underpowered giving our relatively low sample size of 10 splits per cancer. Throughout the text, we used α = 0.05 as the significance level.

We only encountered model failures related to singular fits produced by the *glmnet* R package on certain cancers and methods - as these tended to be systematic, we treated failures as random performance, assigning constant predictions, a concordance of 0.5, and full sparsity (*i*.*e*., no variables selected).

We describe implementational details related to reference methods, students, and teachers in the appendix.

## 5. Results

### 5.1. Knowledge distillation was effective for sparsifying high-dimensional survival models when guided by clinical variables

We found that, when performing knowledge distillation as outlined, standard *𝓁*_1_-regularized linear students (Equation (2)) were able to match their teachers well, achieving reasonably high train and test fidelity as measured by Spearman’s correlation coefficient ρ between the teacher predictions and the student predictions (Table 1). However, in terms of prediction accuracy, only the student of HSAE did not perform significantly worse than its teacher across datasets, with the students of BF and RBF both performing significantly worse than their teachers. All student models were reasonably sparse, with the students of BF and RBF both using an average of roughly 50 features, while the student of HSAE was considerably less sparse at around 130 features.

**Table 1.** Teacher and student discriminative performance, student-teacher prediction rank-correlation and sparsity when pooling across the 17 considered TCGA cancer datasets. Performance of students that did not perform significantly worse (p *≥* 0.05) than their teachers bolded. All p-values based on Wilcoxon ranked sign tests. HSAE: HierarchicalSAE. RBF: RandomBlock favoring. BF: BlockForest.

| TEACHER | STUDENT | MEAN CONCORDANCE | MEAN TRAIN $\rho$ | MEAN TEST $\rho$ | MEAN # FEATURES USED |
| --- | --- | --- | --- | --- | --- |
| HSAE | - | $0.665 \pm 0.104$ | - | - | $79329.8 \pm 3815.2$ |
| RBF | - | $0.663 \pm 0.113$ | - | - | $79329.8 \pm 3815.2$ |
| BF | - | $0.650 \pm 0.118$ | - | - | $79329.8 \pm 3815.2$ |
| HSAE | LASSO | **$0.648 \pm 0.125$** | $0.903 \pm 0.130$ | $0.722 \pm 0.239$ | $131.9 \pm 114.1$ |
| RBF | LASSO | $0.645 \pm 0.120$ | $0.789 \pm 0.123$ | $0.663 \pm 0.212$ | $48.1 \pm 38.8$ |
| BF | LASSO | $0.634 \pm 0.123$ | $0.786 \pm 0.123$ | $0.677 \pm 0.194$ | $51.0 \pm 41.2$ |
| HSAE | LASSO FAVORING | **$0.662 \pm 0.115$** | $0.909 \pm 0.111$ | $0.796 \pm 0.169$ | $139.4 \pm 114.5$ |
| RBF | LASSO FAVORING | **$0.661 \pm 0.117$** | $0.776 \pm 0.121$ | $0.780 \pm 0.125$ | $47.0 \pm 29.1$ |
| BF | LASSO FAVORING | **$0.661 \pm 0.118$** | $0.748 \pm 0.145$ | $0.667 \pm 0.213$ | $51.2 \pm 29.5$ |

Once we enabled unpenalized clinical variables to guide the knowledge distillation (Equation (3)), we found that our sparse linear students were able to learn from their teachers more effectively, with all students except that of BF achieving higher test fidelity as measured by Spearman’s ρ (Table 1), compared to the students which penalized clinical variables. In addition, all students that did not penalize clinical variables achieved discriminative performance very close to that of their teachers (in fact, the student of BF even outperformed its teacher), as none of the clinically guided students performed significantly worse than their teacher. In addition, the clinically guided students achieved sparsity levels comparable with that of the non-clinically guided students (Table 1). Of course, both types of students were orders of magnitude more sparse than their teachers, which did not perform explicit feature selection and thus used all available features (Table 1).

Various factors had an impact on the discriminative performance of our clinically guided students, relative to their teachers. First and foremost, we saw the clinically guided students outperform their teachers on datasets on which the clinical-only model performed particularly well (*e*.*g*., COAD, OV, and LUSC) (Figure 1). However, we also found that our clinically guided students tended to slightly outperform their teachers even on cancers on which the clinical-only model underperformed, such as LGG or LIHC. Overall, we rarely found the clinically guided students to match their teachers exactly in terms of discriminative performance, instead often but not always underperforming (*e*.*g*., SARC) or outperforming them (*e*.*g*., COAD, OV, LGG). Students that penalized clinical variables tended to underperform both their teachers and the clinical-only model on cancers on which the clinical-only model performed particularly well (*e*.*g*., COAD, OV), which was likely part of the reason for their overall underperformance, relative to their teachers (Figure S1).

**Figure 1.**
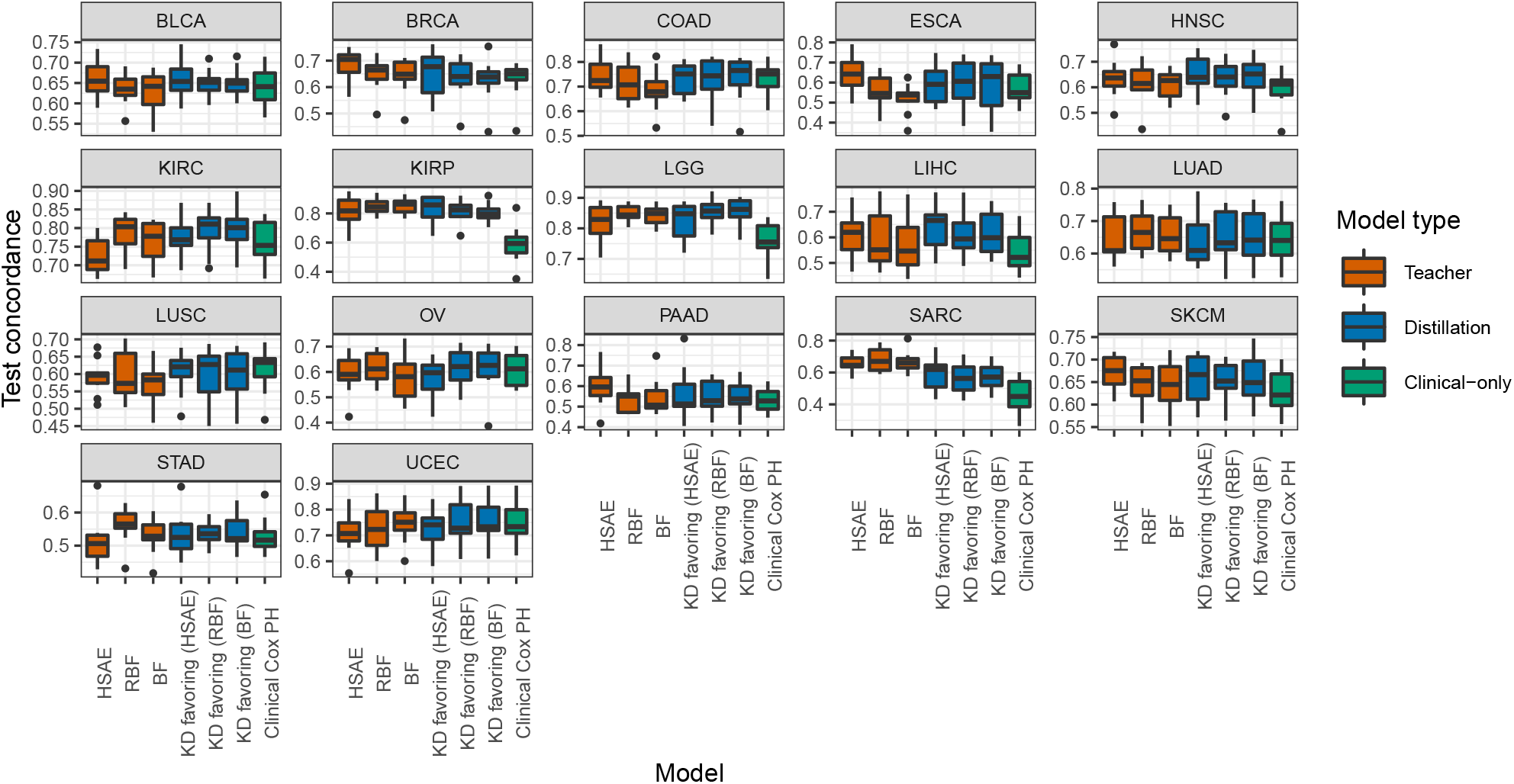
Test concordance per cancer of all teachers and clinically guided students (Equation (3)). Clinical-only model included for reference. HSAE: HierarchicalSAE. RBF: RandomBlock favoring. BF: BlockForest.

We did not find a clear link between student fidelity as measured by Spearman’s ρ and the gap between teacher and student performance. On COAD for example, where all students outperformed their teachers in terms of median test concordance (Figure 1), student fidelity was almost as high as on BLCA (Figure S2), a dataset on which teachers and students performed almost identically in terms of test concordance.

Further, for comparisons to selected reference methods, we restricted our attention to clinically guided students (Equation (3)) due to their improved test fidelity and discriminative performance. Throughout the remainder of the text, we referred to our clinically guided students as knowledge distillation (KD) favoring.

Having established that knowledge distillation could be an effective tool for sparsifying black-box multi-omics survival models, we now move on to concrete comparisons to sparse linear multi-omics survival models from the literature.

### 5.2. Sparse linear students outperformed sparse linear models trained on survival targets

Our student models consistently outperformed the considered sparse reference methods, in addition to the clinical-only Cox PH model (Figure 2 and Figure S3). None of the reference methods was able to outperform the clinical-only Cox PH model significantly as measured by a Wilcoxon ranked sign test across datasets, a feat that all of our students achieved. In addition, our student models achieved similar (for the RBF and BF teachers) or somewhat higher (for the HSAE teacher) sparsity levels across modalities (Table 2), compared to the reference models. Further, we noticed that while *prioritylasso* selected modalities somewhat uniformly, resulting in similar sparsity levels across modalities, our students seemed to focus much more on important modalities such as gene expression. With Lasso favoring, a pattern similar to our students was visible, with a lower sparsity level for seemingly important modalities and higher sparsity for modalities with either lower-dimensional input (*e*.*g*., miRNA) or less predictive value (*e*.*g*., CNV).

**Figure 2.**
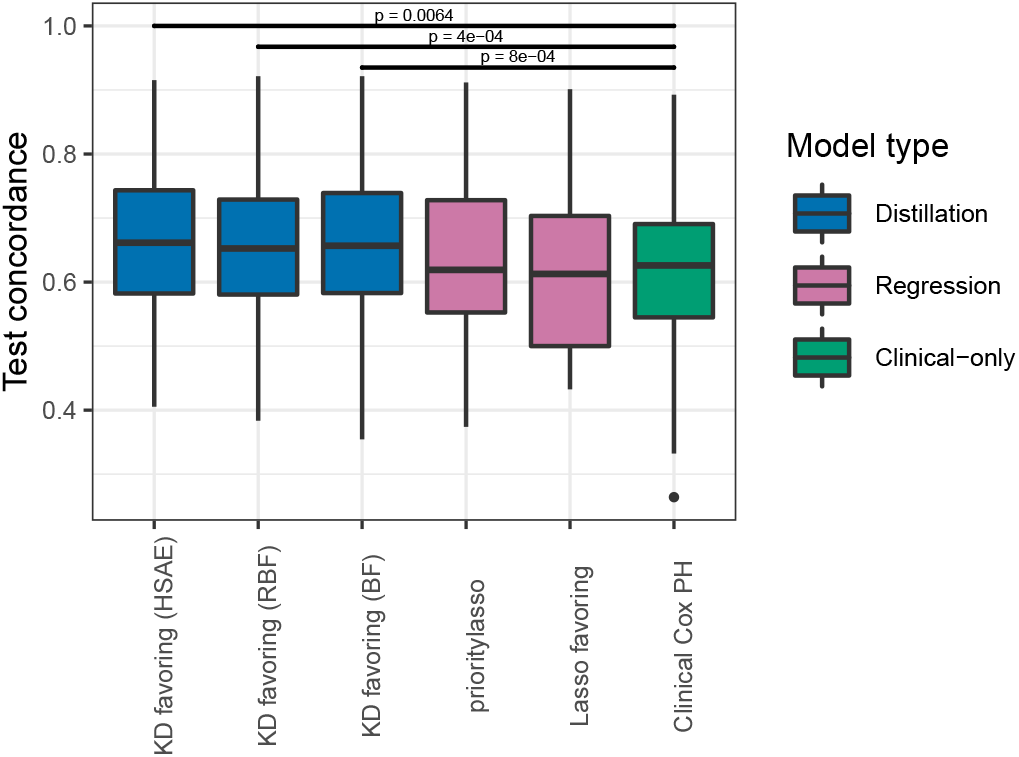
Pooled test concordance of student models and sparse reference models. Clinical-only model included for reference. Significance bars shown for models that outperformed the clinical-only model significantly (p *<* 0.05). All p-values based on Wilcoxon ranked sign tests. HSAE: HierarchicalSAE. RBF: RandomBlock favoring. BF: BlockForest.

**Table 2.**
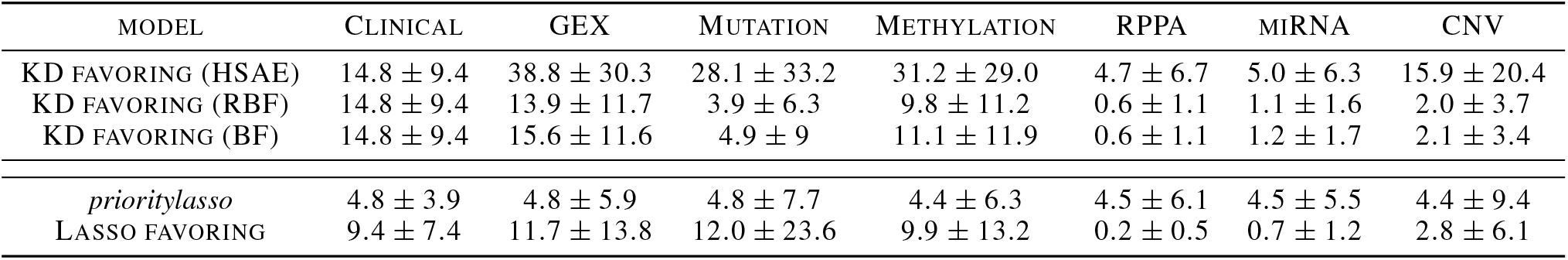
Mean number of selected features in each modality pooled across test splits and cancer types of the proposed students and reference methods. Lasso favoring had a lower number of clinical features due to singular fits which sometimes resulted in empty models. HSAE: HierarchicalSAE. RBF: RandomBlock favoring. BF: BlockForest.

### 5.3. Cross-modal distillation enabled enforced group sparsity while maintaining high performance

While achieving moderate overall sparsity was already an important first step toward enhanced clinical applicability, most clinical settings also require group sparsity, that is sparsity on the level of modalities, both for reasons of costs and interpretability. To accommodate group sparsity, we fitted our student models using cross-modal distillation (Gou et al., 2021). We used the predicted risk measures 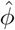 given by teachers trained on the full multi-omics data and distilled their knowledge to students only using a set of two input modalities. We considered clinical data in addition to each available genomic input modality. Reference methods were fit on the same modalities as the students but were trained using the survival targets instead of the teacher knowledge.

Across all modalities, we again saw our students outperform the reference models consistently (Table 3). One of the students performed best in terms of pooled mean concordance in all settings and the reference methods performed significantly worse than the best student in all settings. Further, while all of our students performed significantly better than the clinical-only Cox PH model in at least three settings, the reference methods did not significantly outperform the clinical-only model in any setting. Moreover, the reference methods even performed worse than the clinical only baseline in terms of pooled mean concordance when using clinical and copy number variation or clinical and mutation (Table 3).

**Table 3.**
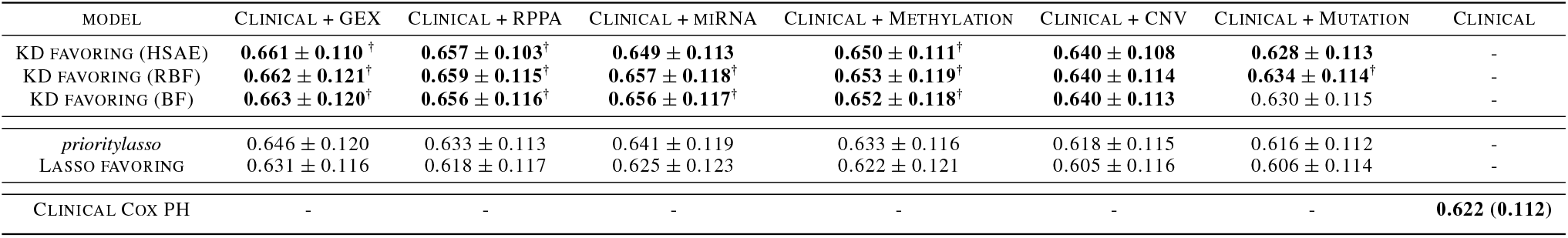
Mean test concordance pooled across datasets of cross-modality distilled students and sparse reference methods when limited to clinical data and each molecular modality. Values not significantly worse (p *≥*0.05) than highest pooled mean concordance bolded by column. Values significantly better (p *<* 0.05) than clinical-only denoted with †. All p-values based on Wilcoxon ranked sign tests. HSAE: HierarchicalSAE. RBF: RandomBlock favoring. BF: BlockForest.

Our results also indicated the heterogeneous predictive value contained within different modalities. Both the students and the reference methods performed much better with some modalities in addition to clinical compared to others. It is well known that some modalities such as gene expression are more useful than others for survival prediction, which has led many models to perform better with only clinical data and gene expression, as opposed to using a larger number of modalities (Hornung & Wright, 2019b; Wissel et al., 2022; Vale-Silva & Rohr, 2021).

### 5.4. Why did clinically guided knowledge distillation outperform the reference models?

Lastly, we were interested in investigating why knowledge distillation had been effective within our context of multiomics integration. Since Lasso favoring had not been able to match the good performance of our students, we had already excluded the possibility that the favoring of clinical variables was the cause of the students’ outperformance (Figure 2). Thus, we next explored to what extent the features selected by our students were responsible for their good performance. To measure the inherent predictive value contained in the features selected by our students and the considered sparse reference methods, we refitted a Ridge Cox PH model using the same feature selected by each sparse method on the survival targets, following the approach of Bommert et al. (2021). If our students or one of the other methods had worked primarily through feature selection, we would have expected for these Ridge models to recover (close to) the same performance as the model that had selected the features.

We found that the two reference methods, *prioritylasso* and Lasso favoring, seemed to perform well primarily due to feature selection, as for both of them, refitting a Ridge Cox PH model with the same feature set resulted in close to the same pooled mean concordance as the original model, even slightly higher for Lasso favoring (Table 4, Δ Originally trained model). That said, both Ridge Cox PH models using the feature sets of the reference methods slightly outperformed the Clinical Cox PH model in terms of pooled mean concordance, although the difference was not significant (Δ Clinical-only). For our clinically guided students, the results were more nuanced. All student models performed better than the Ridge Cox PH models refit with the same feature set, in the cases of BF and HSAE significantly so (Table 4, Δ Originally trained model). The Ridge Cox PH models using the feature sets selected by each of the students also outperformed the clinical-only model in terms of pooled mean concordance, but none of the improvements was significant. Thus, while feature selection seemed to explain part of the good performance of our students, it did not fully account for it, especially for BF and HSAE. Thus, the distillation process did not seem to have worked only through a simple mechanism of feature selection. We leave it to future work to explore the potential existence of other mechanisms contributing to the success of knowledge distillation in survival settings.

**Table 4.** Performance of Ridge Cox PH models fitted with the feature sets selected by each considered sparse method. Δ Clinical-only denotes the mean difference in concordance to the clinical-only Cox PH model. Δ Originally trained model denotes the mean difference in concordance to the originally trained model (that is, the same model that the feature set came from). Models that did not perform significantly worse (p *≥* 0.05) than the model with the highest pooled mean concordance bolded in second column. † denotes significantly (p *<* 0.05) worse performance than the originally trained model that selected the feature set. All p-values based on Wilcoxon ranked sign tests. HSAE: HierarchicalSAE. RBF: RandomBlock favoring. BF: BlockForest.

| FEATURE SET FROM | MEAN TEST CONCORDANCE | $\Delta$ CLINICAL-ONLY | $\Delta$ ORIGINALLY TRAINED MODEL |
| --- | --- | --- | --- |
| KD FAVORING (HSAE) | $0.639 \pm 0.111$ | $0.016 \pm 0.126$ | $-0.024 \pm 0.080^\dagger$ |
| KD FAVORING (RBF) | **$0.654 \pm 0.110$** | $0.032 \pm 0.104$ | $-0.007 \pm 0.057$ |
| KD FAVORING (BF) | $0.647 \pm 0.111$ | $0.025 \pm 0.105$ | $-0.014 \pm 0.058^\dagger$ |
| *prioritylasso* | $0.637 \pm 0.119$ | $0.015 \pm 0.121$ | $0.000 \pm 0.080$ |
| LASSO FAVORING | $0.628 \pm 0.119$ | $0.006 \pm 0.110$ | $0.001 \pm 0.050$ |

## 6. Conclusion

Our work has shown that knowledge distillation can in principle be effectively applied to survival models more broadly, and for the sparsification of black-box multi-omics models on TCGA data more specifically. Moreover, our results suggested that building sparse linear survival models for multi-omics integration could be improved by sparsifying already well-performing black-box models, as opposed to attempting to fit sparse linear models from scratch.

We acknowledge several limitations of our work. First, we limited ourselves to a subset of the TCGA datasets, meaning that our results cannot be guaranteed to generalize to other cancers or databases. In addition, the teachers considered in our work had previously been shown to perform particularly well on TCGA. Further, we restricted our attention to non-pan-cancer models, that is, models that did not leverage transfer learning across multiple cancer types, a limitation that future work should certainly address (Cheerla & Gevaert, 2019; Vale-Silva & Rohr, 2021). Lastly, we restricted ourselves to some student models that could only predict risk measures, limiting their applications to risk stratification in clinical settings. Future work should also investigate the suitability of clinically guided distillation to provide estimates of the survival function.^4^

Future work is further needed in three main areas. First, it is crucial to gain an improved understanding of why knowledge distillation could work in the survival context, beyond the multi-omics setting. Second, one needs to further explore sparse models for multi-omics integration in survival settings, perhaps with a stronger focus on improving performance through proper feature selection. At last, a promising direction for future work is to further explore alternative student models for (clinically guided) knowledge distillation, such as group lasso regularized students.

## Appendix

### A. Implementation

We implemented all teachers using the original code available from the authors, including the choice of hyperparameters. HSAE was implemented using Pytorch (Paszke et al., 2019) and skorch (Tietz et al., 2017), while BF and RBF were implemented using the *BlockForest* R package written by the authors (Hornung & Wright, 2019a). All student models were implemented using the *glmnet* R package (Friedman et al., 2010). For the students, we used the regularization hyperparameter λ that minimized the mean squared error between teacher and student predictions using standard five-fold cross-validation (without stratifying for the censoring indicator).

Our two reference models, *prioritylasso* and Lasso favoring were implemented using the *prioritylasso* R package (Klau et al., 2020) and the *glmnet* R package respectively. For both reference models, we used the regularization hyperparameter λ that minimized the negative partial log-likelihood using standard five-fold cross-validation (without stratifying for the censoring indicator).

We pre-specified the priority order of *prioritylasso* to be clinical, gene expression, mutation, miRNA, DNA methylation, CNV, and RPPA, in descending order of importance. Although clinicians and users may not know the ideal priority order on an arbitrary dataset, we wanted to give *prioritylasso* a chance to perform at its best, thus giving it the advantage of specifying a priority order which was likely better than something the model would have been able to learn on its own (as the authors in Herrmann et al. (2021) did). Gene expression is known to contain ample information on TCGA, with multi-omics models often performing better when only using clinical data and gene expression as opposed to multiple modalities (Vale-Silva & Rohr, 2021; Hornung & Wright, 2019b; Wissel et al., 2022). Although our given priority ordering was likely still not optimal for all cancers, at least choosing clinical and gene expression as the two most important modalities already gave the model an advantage. We initially did not penalize clinical variables for *prioritylasso* (deemed *prioritylasso favoring* (Klau et al., 2018; Herrmann et al., 2021)), but found that this sometimes caused convergence issues and did not notably improve performance as long as the model was given a decent priority order. Thus, we fitted the clinical modality *𝓁*_1_-penalized, like all other modalities.

All random seeds were fixed to 42. Our full implementation is available as a supplement to this submission.

### B. Supplementary figures

**Figure S1.**
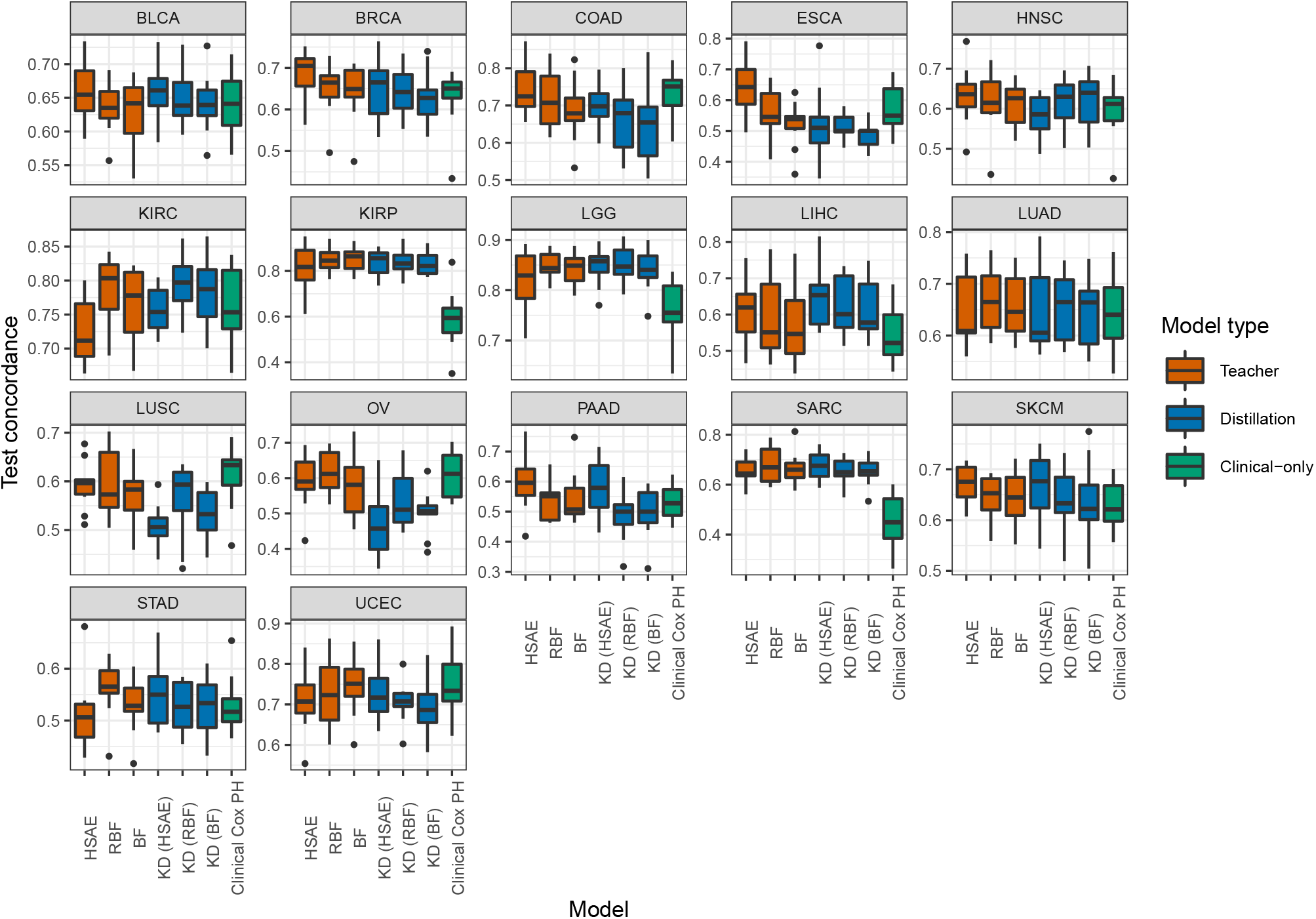
Test concordance of teachers and students that penalized clinical variables (Equation (2)) per cancer. We found that the overall underperformance of students penalizing clinical variables was driven by cancers such as LUSC, COAD and OV, on which the students penalizing clinical variables underperformed both the clinical-only model and their teachers. HSAE: HierarchicalSAE. RBF: RandomBlock favoring. BF: BlockForest.

**Figure S2.**
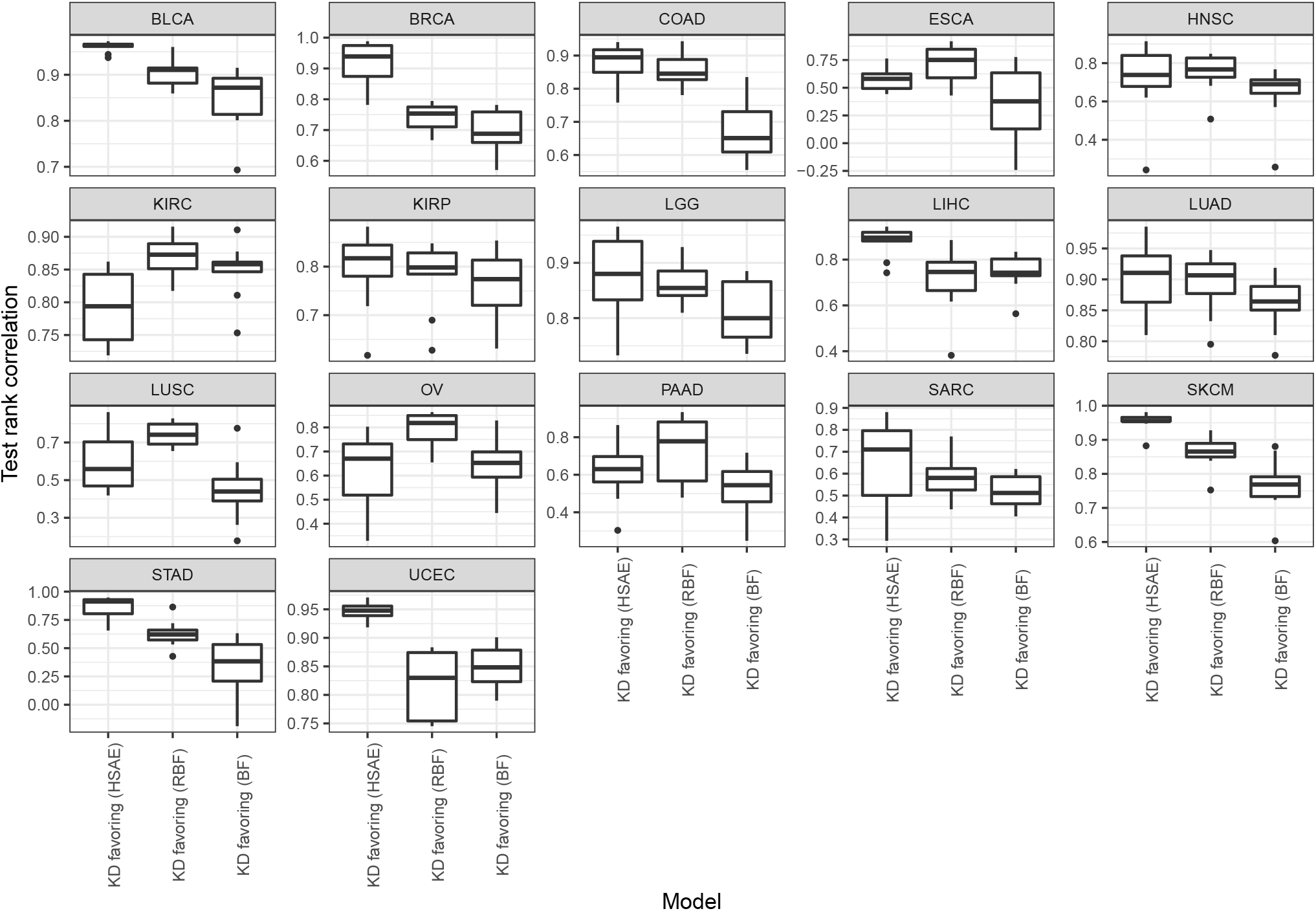
Spearman’s *ρ* between the teacher and clinically guided student test predictions across datasets. While we observed large hetereogeneity in the generalization gap between teacher and student within our knowledge distillation, the size or direction of this gap was seemingly not directly related to student fidelity in terms of Spearman’s *ρ* on the test set. HSAE: HierarchicalSAE. RBF: RandomBlock favoring. BF: BlockForest.

**Figure S3.**
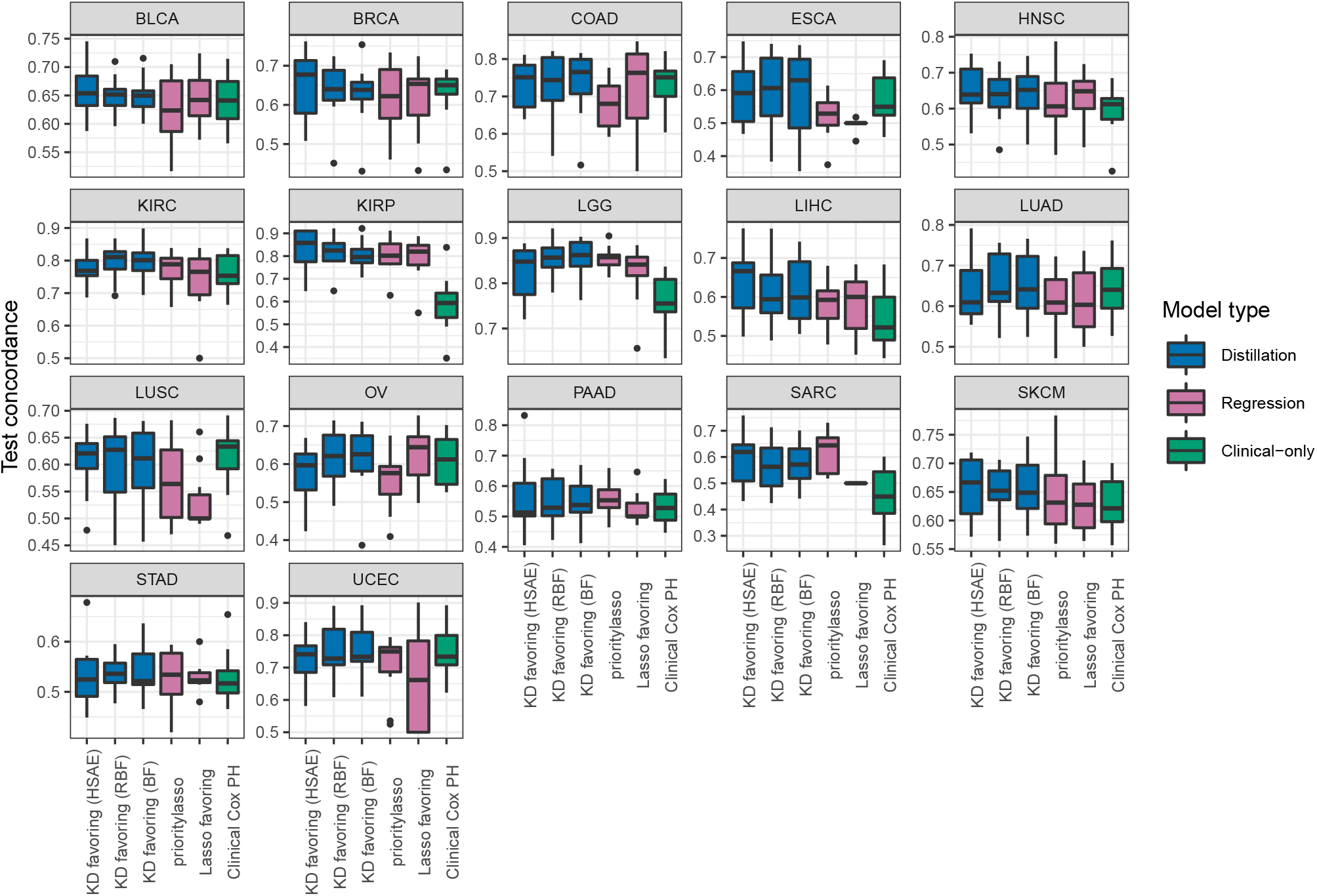
Test concordance per cancer of student models and sparse reference models. Clinical-only model included for reference. We saw our students outperform both the clinical-only model and the reference methods for most cancers. HSAE: HierarchicalSAE. RBF: RandomBlock favoring. BF: BlockForest.

## Footnotes

1 Throughout our work, we use the term *risk measure* loosely to refer to different predicted outcomes of survival models which all share in common that if patient *i* has a higher predicted risk measure than patient *j*, patient *i* is expected (according to the model) to die before *j*.

2 Presumably, using fewer clinical variables relative to Herrmann et al. (2021) could be expected to reduce the performance of the clinical-only model.

3 For random forest methods, since we used ensemble mortality as the response-based knowledge, there was no simple way for the students to produce estimates of the survival function.

4 Using Cox PH-based log-risk as the response-based knowledge provided by the teacher, for example, could enable the students to produce estimates of the survival function as well.

