## Supplementary material for "Clinically driven knowledge distillation for sparsifying high-dimensional multi-omics survival models": Code: README.md

# Clinically driven knowledge distillation for sparsifying high-dimensional multi-omics survival models
## Reproduction guide
To reproduce any of our results, all you have to do is run the requisite bash scripts. 

Our preprocessed data can be downloaded [here](https://drive.google.com/file/d/1K7dexVJhe-vVoQ0XeKh6YtTyfu4T_jZe/view).

Please make sure that you have [R](https://www.r-project.org/) 4.0.X and a current version of [conda](https://docs.conda.io/en/latest/miniconda.html) installed before attempting to reproduce any of the below. You also need a `Fortran` compiler to use `glmnet` - if you don't have one already, you can install it with `brew install gcc` if you have `brew` already. 

If you do not have `R` 4.0.X (but an older version), we currently handle this by force installing all packages (without using `renv`) - note that this may fail (if one of the packages is not available for your `R` version) or produce unexpected results (in case one of the packages is not forward- or backward-compatible with the version that we were using). Unfortunately we did not find an easy way to install a specific `R` version outside of using a Docker container.

### Setup
`bash setup.sh`

Please note that you might have to run the setup script with `sudo` depending on your setup.

### Re-running benchmarks
To re-run all benchmarks, run `bash reproduce_benchmarks.sh`. Of course, you can also just re-run single scripts manually. Note that some scripts pre-suppose the existance of certain files (e.g., you cannot run the distillation before you have run the teachers). For the "correct" order, simply have a look at the `reproduce_benchmarks.sh` script (in case you want to run individual scripts).

Please note that in order to reproduce figures and tables, you need to first re-run the benchmarks.

### Reproducing figures
`bash reproduce_figures.sh`

### Reproducing tables

`bash reproduce_tables.sh`
