## Supplementary material for "Clinically driven knowledge distillation for sparsifying high-dimensional multi-omics survival models": Code: activate.R

```
local({

  # the requested version of renv
  version <- "0.14.0"

  # the project directory
  project <- getwd()

  # allow environment variable to control activation
  activate <- Sys.getenv("RENV_ACTIVATE_PROJECT")
  if (!nzchar(activate)) {

    # don't auto-activate when R CMD INSTALL is running
    if (nzchar(Sys.getenv("R_INSTALL_PKG")))
      return(FALSE)

  }

  # bail if activation was explicitly disabled
  if (tolower(activate) %in% c("false", "f", "0"))
    return(FALSE)

  # avoid recursion
  if (nzchar(Sys.getenv("RENV_R_INITIALIZING")))
    return(invisible(TRUE))

  # signal that we're loading renv during R startup
  Sys.setenv("RENV_R_INITIALIZING" = "true")
  on.exit(Sys.unsetenv("RENV_R_INITIALIZING"), add = TRUE)

  # signal that we've consented to use renv
  options(renv.consent = TRUE)

  # load the 'utils' package eagerly -- this ensures that renv shims, which
  # mask 'utils' packages, will come first on the search path
  library(utils, lib.loc = .Library)

  # check to see if renv has already been loaded
  if ("renv" %in% loadedNamespaces()) {

    # if renv has already been loaded, and it's the requested version of renv,
    # nothing to do
    spec <- .getNamespaceInfo(.getNamespace("renv"), "spec")
    if (identical(spec[["version"]], version))
      return(invisible(TRUE))

    # otherwise, unload and attempt to load the correct version of renv
    unloadNamespace("renv")

  }

  # load bootstrap tools   
  bootstrap <- function(version, library) {
  
    # attempt to download renv
    tarball <- tryCatch(renv_bootstrap_download(version), error = identity)
    if (inherits(tarball, "error"))
      stop("failed to download renv ", version)
  
    # now attempt to install
    status <- tryCatch(renv_bootstrap_install(version, tarball, library), error = identity)
    if (inherits(status, "error"))
      stop("failed to install renv ", version)
  
  }
  
  renv_bootstrap_tests_running <- function() {
    getOption("renv.tests.running", default = FALSE)
  }
  
  renv_bootstrap_repos <- function() {
  
    # check for repos override
    repos <- Sys.getenv("RENV_CONFIG_REPOS_OVERRIDE", unset = NA)
    if (!is.na(repos))
      return(repos)
  
    # if we're testing, re-use the test repositories
    if (renv_bootstrap_tests_running())
      return(getOption("renv.tests.repos"))
  
    # retrieve current repos
    repos <- getOption("repos")
  
    # ensure @CRAN@ entries are resolved
    repos[repos == "@CRAN@"] <- getOption(
      "renv.repos.cran",
      "https://cloud.r-project.org"
    )
  
    # add in renv.bootstrap.repos if set
    default <- c(FALLBACK = "https://cloud.r-project.org")
    extra <- getOption("renv.bootstrap.repos", default = default)
    repos <- c(repos, extra)
  
    # remove duplicates that might've snuck in
    dupes <- duplicated(repos) | duplicated(names(repos))
    repos[!dupes]
  
  }
  
  renv_bootstrap_download <- function(version) {
  
    # if the renv version number has 4 components, assume it must
    # be retrieved via github
    nv <- numeric_version(version)
    components <- unclass(nv)[[1]]
  
    methods <- if (length(components) == 4L) {
      list(
        renv_bootstrap_download_github
      )
    } else {
      list(
        renv_bootstrap_download_cran_latest,
        renv_bootstrap_download_cran_archive
      )
    }
  
    for (method in methods) {
      path <- tryCatch(method(version), error = identity)
      if (is.character(path) && file.exists(path))
        return(path)
    }
  
    stop("failed to download renv ", version)
  
  }
  
  renv_bootstrap_download_impl <- function(url, destfile) {
  
    mode <- "wb"
  
    # https://bugs.r-project.org/bugzilla/show_bug.cgi?id=17715
    fixup <-
      Sys.info()[["sysname"]] == "Windows" &&
      substring(url, 1L, 5L) == "file:"
  
    if (fixup)
      mode <- "w+b"
  
    utils::download.file(
      url      = url,
      destfile = destfile,
      mode     = mode,
      quiet    = TRUE
    )
  
  }
  
  renv_bootstrap_download_cran_latest <- function(version) {
  
    spec <- renv_bootstrap_download_cran_latest_find(version)
  
    message("* Downloading renv ", version, " ... ", appendLF = FALSE)
  
    type  <- spec$type
    repos <- spec$repos
  
    info <- tryCatch(
      utils::download.packages(
        pkgs    = "renv",
        destdir = tempdir(),
        repos   = repos,
        type    = type,
        quiet   = TRUE
      ),
      condition = identity
    )
  
    if (inherits(info, "condition")) {
      message("FAILED")
      return(FALSE)
    }
  
    # report success and return
    message("OK (downloaded ", type, ")")
    info[1, 2]
  
  }
  
  renv_bootstrap_download_cran_latest_find <- function(version) {
  
    # check whether binaries are supported on this system
    binary <-
      getOption("renv.bootstrap.binary", default = TRUE) &&
      !identical(.Platform$pkgType, "source") &&
      !identical(getOption("pkgType"), "source") &&
      Sys.info()[["sysname"]] %in% c("Darwin", "Windows")
  
    types <- c(if (binary) "binary", "source")
  
    # iterate over types + repositories
    for (type in types) {
      for (repos in renv_bootstrap_repos()) {
  
        # retrieve package database
        db <- tryCatch(
          as.data.frame(
            utils::available.packages(type = type, repos = repos),
            stringsAsFactors = FALSE
          ),
          error = identity
        )
  
        if (inherits(db, "error"))
          next
  
        # check for compatible entry
        entry <- db[db$Package %in% "renv" & db$Version %in% version, ]
        if (nrow(entry) == 0)
          next
  
        # found it; return spec to caller
        spec <- list(entry = entry, type = type, repos = repos)
        return(spec)
  
      }
    }
  
    # if we got here, we failed to find renv
    fmt <- "renv %s is not available from your declared package repositories"
    stop(sprintf(fmt, version))
  
  }
  
  renv_bootstrap_download_cran_archive <- function(version) {
  
    name <- sprintf("renv_%s.tar.gz", version)
    repos <- renv_bootstrap_repos()
    urls <- file.path(repos, "src/contrib/Archive/renv", name)
    destfile <- file.path(tempdir(), name)
  
    message("* Downloading renv ", version, " ... ", appendLF = FALSE)
  
    for (url in urls) {
  
      status <- tryCatch(
        renv_bootstrap_download_impl(url, destfile),
        condition = identity
      )
  
      if (identical(status, 0L)) {
        message("OK")
        return(destfile)
      }
  
    }
  
    message("FAILED")
    return(FALSE)
  
  }
  
  renv_bootstrap_download_github <- function(version) {
  
    enabled <- Sys.getenv("RENV_BOOTSTRAP_FROM_GITHUB", unset = "TRUE")
    if (!identical(enabled, "TRUE"))
      return(FALSE)
  
    # prepare download options
    pat <- Sys.getenv("GITHUB_PAT")
    if (nzchar(Sys.which("curl")) && nzchar(pat)) {
      fmt <- "--location --fail --header \"Authorization: token %s\""
      extra <- sprintf(fmt, pat)
      saved <- options("download.file.method", "download.file.extra")
      options(download.file.method = "curl", download.file.extra = extra)
      on.exit(do.call(base::options, saved), add = TRUE)
    } else if (nzchar(Sys.which("wget")) && nzchar(pat)) {
      fmt <- "--header=\"Authorization: token %s\""
      extra <- sprintf(fmt, pat)
      saved <- options("download.file.method", "download.file.extra")
      options(download.file.method = "wget", download.file.extra = extra)
      on.exit(do.call(base::options, saved), add = TRUE)
    }
  
    message("* Downloading renv ", version, " from GitHub ... ", appendLF = FALSE)
  
    url <- file.path("https://api.github.com/repos/rstudio/renv/tarball", version)
    name <- sprintf("renv_%s.tar.gz", version)
    destfile <- file.path(tempdir(), name)
  
    status <- tryCatch(
      renv_bootstrap_download_impl(url, destfile),
      condition = identity
    )
  
    if (!identical(status, 0L)) {
      message("FAILED")
      return(FALSE)
    }
  
    message("OK")
    return(destfile)
  
  }
  
  renv_bootstrap_install <- function(version, tarball, library) {
  
    # attempt to install it into project library
    message("* Installing renv ", version, " ... ", appendLF = FALSE)
    dir.create(library, showWarnings = FALSE, recursive = TRUE)
  
    # invoke using system2 so we can capture and report output
    bin <- R.home("bin")
    exe <- if (Sys.info()[["sysname"]] == "Windows") "R.exe" else "R"
    r <- file.path(bin, exe)
    args <- c("--vanilla", "CMD", "INSTALL", "-l", shQuote(library), shQuote(tarball))
    output <- system2(r, args, stdout = TRUE, stderr = TRUE)
    message("Done!")
  
    # check for successful install
    status <- attr(output, "status")
    if (is.numeric(status) && !identical(status, 0L)) {
      header <- "Error installing renv:"
      lines <- paste(rep.int("=", nchar(header)), collapse = "")
      text <- c(header, lines, output)
      writeLines(text, con = stderr())
    }
  
    status
  
  }
  
  renv_bootstrap_platform_prefix <- function() {
  
    # construct version prefix
    version <- paste(R.version$major, R.version$minor, sep = ".")
    prefix <- paste("R", numeric_version(version)[1, 1:2], sep = "-")
  
    # include SVN revision for development versions of R
    # (to avoid sharing platform-specific artefacts with released versions of R)
    devel <-
      identical(R.version[["status"]],   "Under development (unstable)") ||
      identical(R.version[["nickname"]], "Unsuffered Consequences")
  
    if (devel)
      prefix <- paste(prefix, R.version[["svn rev"]], sep = "-r")
  
    # build list of path components
    components <- c(prefix, R.version$platform)
  
    # include prefix if provided by user
    prefix <- renv_bootstrap_platform_prefix_impl()
    if (!is.na(prefix) && nzchar(prefix))
      components <- c(prefix, components)
  
    # build prefix
    paste(components, collapse = "/")
  
  }
  
  renv_bootstrap_platform_prefix_impl <- function() {
  
    # if an explicit prefix has been supplied, use it
    prefix <- Sys.getenv("RENV_PATHS_PREFIX", unset = NA)
    if (!is.na(prefix))
      return(prefix)
  
    # if the user has requested an automatic prefix, generate it
    auto <- Sys.getenv("RENV_PATHS_PREFIX_AUTO", unset = NA)
    if (auto %in% c("TRUE", "True", "true", "1"))
      return(renv_bootstrap_platform_prefix_auto())
  
    # empty string on failure
    ""
  
  }
  
  renv_bootstrap_platform_prefix_auto <- function() {
  
    prefix <- tryCatch(renv_bootstrap_platform_os(), error = identity)
    if (inherits(prefix, "error") || prefix %in% "unknown") {
  
      msg <- paste(
        "failed to infer current operating system",
        "please file a bug report at https://github.com/rstudio/renv/issues",
        sep = "; "
      )
  
      warning(msg)
  
    }
  
    prefix
  
  }
  
  renv_bootstrap_platform_os <- function() {
  
    sysinfo <- Sys.info()
    sysname <- sysinfo[["sysname"]]
  
    # handle Windows + macOS up front
    if (sysname == "Windows")
      return("windows")
    else if (sysname == "Darwin")
      return("macos")
  
    # check for os-release files
    for (file in c("/etc/os-release", "/usr/lib/os-release"))
      if (file.exists(file))
        return(renv_bootstrap_platform_os_via_os_release(file, sysinfo))
  
    # check for redhat-release files
    if (file.exists("/etc/redhat-release"))
      return(renv_bootstrap_platform_os_via_redhat_release())
  
    "unknown"
  
  }
  
  renv_bootstrap_platform_os_via_os_release <- function(file, sysinfo) {
  
    # read /etc/os-release
    release <- utils::read.table(
      file             = file,
      sep              = "=",
      quote            = c("\"", "'"),
      col.names        = c("Key", "Value"),
      comment.char     = "#",
      stringsAsFactors = FALSE
    )
  
    vars <- as.list(release$Value)
    names(vars) <- release$Key
  
    # get os name
    os <- tolower(sysinfo[["sysname"]])
  
    # read id
    id <- "unknown"
    for (field in c("ID", "ID_LIKE")) {
      if (field %in% names(vars) && nzchar(vars[[field]])) {
        id <- vars[[field]]
        break
      }
    }
  
    # read version
    version <- "unknown"
    for (field in c("UBUNTU_CODENAME", "VERSION_CODENAME", "VERSION_ID", "BUILD_ID")) {
      if (field %in% names(vars) && nzchar(vars[[field]])) {
        version <- vars[[field]]
        break
      }
    }
  
    # join together
    paste(c(os, id, version), collapse = "-")
  
  }
  
  renv_bootstrap_platform_os_via_redhat_release <- function() {
  
    # read /etc/redhat-release
    contents <- readLines("/etc/redhat-release", warn = FALSE)
  
    # infer id
    id <- if (grepl("centos", contents, ignore.case = TRUE))
      "centos"
    else if (grepl("redhat", contents, ignore.case = TRUE))
      "redhat"
    else
      "unknown"
  
    # try to find a version component (very hacky)
    version <- "unknown"
  
    parts <- strsplit(contents, "[[:space:]]")[[1L]]
    for (part in parts) {
  
      nv <- tryCatch(numeric_version(part), error = identity)
      if (inherits(nv, "error"))
        next
  
      version <- nv[1, 1]
      break
  
    }
  
    paste(c("linux", id, version), collapse = "-")
  
  }
  
  renv_bootstrap_library_root_name <- function(project) {
  
    # use project name as-is if requested
    asis <- Sys.getenv("RENV_PATHS_LIBRARY_ROOT_ASIS", unset = "FALSE")
    if (asis)
      return(basename(project))
  
    # otherwise, disambiguate based on project's path
    id <- substring(renv_bootstrap_hash_text(project), 1L, 8L)
    paste(basename(project), id, sep = "-")
  
  }
  
  renv_bootstrap_library_root <- function(project) {
  
    path <- Sys.getenv("RENV_PATHS_LIBRARY", unset = NA)
    if (!is.na(path))
      return(path)
  
    path <- Sys.getenv("RENV_PATHS_LIBRARY_ROOT", unset = NA)
    if (!is.na(path)) {
      name <- renv_bootstrap_library_root_name(project)
      return(file.path(path, name))
    }
  
    prefix <- renv_bootstrap_profile_prefix()
    paste(c(project, prefix, "renv/library"), collapse = "/")
  
  }
  
  renv_bootstrap_validate_version <- function(version) {
  
    loadedversion <- utils::packageDescription("renv", fields = "Version")
    if (version == loadedversion)
      return(TRUE)
  
    # assume four-component versions are from GitHub; three-component
    # versions are from CRAN
    components <- strsplit(loadedversion, "[.-]")[[1]]
    remote <- if (length(components) == 4L)
      paste("rstudio/renv", loadedversion, sep = "@")
    else
      paste("renv", loadedversion, sep = "@")
  
    fmt <- paste(
      "renv %1$s was loaded from project library, but this project is configured to use renv %2$s.",
      "Use `renv::record(\"%3$s\")` to record renv %1$s in the lockfile.",
      "Use `renv::restore(packages = \"renv\")` to install renv %2$s into the project library.",
      sep = "\n"
    )
  
    msg <- sprintf(fmt, loadedversion, version, remote)
    warning(msg, call. = FALSE)
  
    FALSE
  
  }
  
  renv_bootstrap_hash_text <- function(text) {
  
    hashfile <- tempfile("renv-hash-")
    on.exit(unlink(hashfile), add = TRUE)
  
    writeLines(text, con = hashfile)
    tools::md5sum(hashfile)
  
  }
  
  renv_bootstrap_load <- function(project, libpath, version) {
  
    # try to load renv from the project library
    if (!requireNamespace("renv", lib.loc = libpath, quietly = TRUE))
      return(FALSE)
  
    # warn if the version of renv loaded does not match
    renv_bootstrap_validate_version(version)
  
    # load the project
    renv::load(project)
  
    TRUE
  
  }
  
  renv_bootstrap_profile_load <- function(project) {
  
    # if RENV_PROFILE is already set, just use that
    profile <- Sys.getenv("RENV_PROFILE", unset = NA)
    if (!is.na(profile) && nzchar(profile))
      return(profile)
  
    # check for a profile file (nothing to do if it doesn't exist)
    path <- file.path(project, "renv/local/profile")
    if (!file.exists(path))
      return(NULL)
  
    # read the profile, and set it if it exists
    contents <- readLines(path, warn = FALSE)
    if (length(contents) == 0L)
      return(NULL)
  
    # set RENV_PROFILE
    profile <- contents[[1L]]
    if (nzchar(profile))
      Sys.setenv(RENV_PROFILE = profile)
  
    profile
  
  }
  
  renv_bootstrap_profile_prefix <- function() {
    profile <- renv_bootstrap_profile_get()
    if (!is.null(profile))
      return(file.path("renv/profiles", profile))
  }
  
  renv_bootstrap_profile_get <- function() {
    profile <- Sys.getenv("RENV_PROFILE", unset = "")
    renv_bootstrap_profile_normalize(profile)
  }
  
  renv_bootstrap_profile_set <- function(profile) {
    profile <- renv_bootstrap_profile_normalize(profile)
    if (is.null(profile))
      Sys.unsetenv("RENV_PROFILE")
    else
      Sys.setenv(RENV_PROFILE = profile)
  }
  
  renv_bootstrap_profile_normalize <- function(profile) {
  
    if (is.null(profile) || profile %in% c("", "default"))
      return(NULL)
  
    profile
  
  }

  # load the renv profile, if any
  renv_bootstrap_profile_load(project)

  # construct path to library root
  root <- renv_bootstrap_library_root(project)

  # construct library prefix for platform
  prefix <- renv_bootstrap_platform_prefix()

  # construct full libpath
  libpath <- file.path(root, prefix)

  # attempt to load
  if (renv_bootstrap_load(project, libpath, version))
    return(TRUE)

  # load failed; inform user we're about to bootstrap
  prefix <- paste("# Bootstrapping renv", version)
  postfix <- paste(rep.int("-", 77L - nchar(prefix)), collapse = "")
  header <- paste(prefix, postfix)
  message(header)

  # perform bootstrap
  bootstrap(version, libpath)

  # exit early if we're just testing bootstrap
  if (!is.na(Sys.getenv("RENV_BOOTSTRAP_INSTALL_ONLY", unset = NA)))
    return(TRUE)

  # try again to load
  if (requireNamespace("renv", lib.loc = libpath, quietly = TRUE)) {
    message("* Successfully installed and loaded renv ", version, ".")
    return(renv::load())
  }

  # failed to download or load renv; warn the user
  msg <- c(
    "Failed to find an renv installation: the project will not be loaded.",
    "Use `renv::activate()` to re-initialize the project."
  )

  warning(paste(msg, collapse = "\n"), call. = FALSE)

})
```
