## Supplementary material for "Clinically driven knowledge distillation for sparsifying high-dimensional multi-omics survival models": Code: reproduce_benchmarks.sh

```
#!/bin/bash

Rscript ./src/jobs/run_baselines_main.R
Rscript ./src/jobs/run_random_forest_teachers.R

CONDA_PATH=$(conda info | grep -i 'base environment' | awk '{print $4}')
source $CONDA_PATH/etc/profile.d/conda.sh
conda activate clinical_distillation

python ./src/jobs/run_neural_teacher.py

Rscript ./src/jobs/run_students_main.R
Rscript ./src/jobs/run_students_with_intercept.R

Rscript ./src/jobs/run_table_3_students.R
Rscript ./src/jobs/run_table_3_baselines.R
Rscript ./src/jobs/run_table_4_experiment.R
```
