## Supplementary material for "Clinically driven knowledge distillation for sparsifying high-dimensional multi-omics survival models": Code: setup.R

```
r = getOption("repos")
r["CRAN"] = "https://cloud.r-project.org/"
options(repos = r)
install.packages("renv")
install.packages("here")
library(here)
library(renv)

tryCatch(
  expr = {
    renv::restore(here::here())
  },
  error = function(e){ 
    packages <- readLines(here::here("R_requirements.txt"))
    renv::install("BiocManager")
    library(BiocManager)
    options(repos = BiocManager::repositories())
    renv::install(packages)
  }
)
```
