## Supplementary material for "Clinically driven knowledge distillation for sparsifying high-dimensional multi-omics survival models": Code: setup.sh

```
#!/bin/bash

CONDA_PATH=$(conda info | grep -i 'base environment' | awk '{print $4}')
conda create --name thesis
source $CONDA_PATH/etc/profile.d/conda.sh
conda activate thesis
conda config --add channels conda-forge
conda install pytorch
conda install numpy
conda install scikit-learn
conda install skorch
conda install pandas
conda install -c conda-forge lifelines
pip install glmnet

Rscript setup.R
```
