## Supplementary material for "Clinically driven knowledge distillation for sparsifying high-dimensional multi-omics survival models": Code: run_baselines_main.R

```
library(here)
library(rjson)
library(readr)
library(vroom)
library(dplyr)

source(here::here("src", "utils", "utils.R"))
source(here::here("src", "model", "statistical_models_multi_omics.R"))
config <- rjson::fromJSON(
  file = here::here("config", "config.json")
)

def_order <- c(
  "clinical",
  "gex",
  "mut",
  "mirna",
  "meth",
  "cnv",
  "rppa"
)
modal_mapping <- list(
  "7" = "all"
)

helper_mapping <- list(
  "7" = def_order
)

for (cancer in config$cancers) {
  print(paste0("Starting cancer: ", cancer))
  set.seed(config$seed)
  data <- data.frame(vroom::vroom(
    here::here(
      "data", "processed", cancer, "merged", config$data_name_tcga
    ),
  ), check.names = FALSE)
  train_splits <- read_csv(
    here::here(
      "data", "splits", cancer, config$train_split_name_tcga
    )
  )

  test_splits <- read_csv(
    here::here(
      "data", "splits", cancer, config$test_split_name_tcga
    )
  )

  train_splits <- format_splits(train_splits)
  test_splits <- format_splits(test_splits)
  models <- c("glmnet_favor", "prioritylasso")
  for (model in models) {
    for (modalities in c(7)) {
      print(paste0("Starting model: ", model))
      results <- list()
      for (i in 1:length(train_splits)) {
        print(i)
        train_ix <- train_splits[[i]]
        test_ix <- test_splits[[i]]
        X_train <- data[
            train_ix,
            unname(unlist(sapply(helper_mapping[[as.character(modalities)]], function(block) grep(block, colnames(data)))))
          ] %>%
            data.matrix()
        X_test <- data[
            test_ix,
            unname(
              unlist(sapply(helper_mapping[[as.character(modalities)]], function(block) grep(block, colnames(data))))
            )
          ] %>%
            data.matrix()
    
        y_train <- data.matrix(data[train_ix, 2:1])
        y_test <- data.matrix(data[test_ix, 2:1])
        blocks <- sapply(helper_mapping[[as.character(modalities)]], function(block) grep(block, colnames(X_train)))
        names(blocks) <- paste0(rep("bp", length(helper_mapping[[as.character(modalities)]])), 1:length(helper_mapping[[as.character(modalities)]]))
        penalty_factor <- NA

        tryCatch(
          {
            results[[i]] <- run_benchmark(
              model = model,
              X_train = X_train,
              y_train = y_train,
              X_test = X_test,
              y_test = y_test,
              blocks,
              seed = config$seed
            )
          },
          error = function(cond) {
            # In the case of failure, we assign NA values
            # which we later replace with random performance
            # for concordance and complete sparsity for 
            # the number of features used.
            # Note: Need to make sure to use
            # global assignment here.
            print(cond)
            print("Error running model.")
            print("Saving everything as NA.")
            results[[i]] <<- list(
              concordance = NA,
              ibs = NA,
              params = NA,
              selected_variables = NA,
              train_predictions = NA,
              test_predictions = NA
            )
          }
        )
      }
      saveRDS(
        results,
        file = here::here("data", "benchmarks", cancer, paste0(model, paste0("_10_stratified_", modal_mapping[as.character(modalities)], ".rds")))
      )
    }
  }
}
```
