## Supplementary material for "Clinically driven knowledge distillation for sparsifying high-dimensional multi-omics survival models": Code: run_neural_teacher.py

```
#!/usr/bin/env python
# -*- coding: utf-8 -*-


import json
import os
import sys

module_path = os.path.abspath(os.path.join("./src/"))
if module_path not in sys.path:
    sys.path.append(module_path)

import numpy as np
import pandas as pd
import torch
from model.autoencoders import HierarchicalSAE, HierarchicalSAENet
from sklearn.pipeline import make_pipeline
from sklearn.preprocessing import StandardScaler
from skorch.callbacks import EarlyStopping, LRScheduler
from utils.utils import (
    FixRandomSeed,
    StratifiedSkorchSurvivalSplit,
    hierarchical_sae_criterion,
)

# All code adapted from:
# Wissel, D., Rowson, D., and Boeva, V. Hierarchical autoencoder-based integration improves performance in multi-omics cancer survival models through soft modality selection. bioRxiv, 2022.
def main():
    with open("config/config.json") as f:
        config = json.load(f)
    model = "hierarchicalsaenet"
    modal_mapping = {
        7: "all",
    }
    for cancer in config["cancers"]:
        data = pd.read_csv(
            f"./data/processed/{cancer}/merged/{config['data_name_tcga']}"
        )
        for modalities in [7]:
            print(f"Starting: {cancer}")
            X = data[data.columns[2:]]
            X = X.loc[:, (X != X.iloc[0]).any()]
            y_str = data["OS"].astype(str) + "|" + data["OS.time"].astype(str)

            train_splits = pd.read_csv(
                f"./data/splits/{cancer}/{config['train_split_name_tcga']}"
            )
            test_splits = pd.read_csv(
                f"./data/splits/{cancer}/{config['test_split_name_tcga']}"
            )
            clinical_indices = [
                i for i in range(len(X.columns)) if "clinical" in X.columns[i]
            ]
            gex_indices = [
                i for i in range(len(X.columns)) if "gex" in X.columns[i]
            ]
            cnv_indices = [
                i for i in range(len(X.columns)) if "cnv" in X.columns[i]
            ]
            meth_indices = [
                i for i in range(len(X.columns)) if "meth" in X.columns[i]
            ]
            mirna_indices = [
                i for i in range(len(X.columns)) if "mirna" in X.columns[i]
            ]
            mut_indices = [
                i for i in range(len(X.columns)) if "mut" in X.columns[i]
            ]
            rppa_indices = [
                i for i in range(len(X.columns)) if "rppa" in X.columns[i]
            ]

            blocks = [
                clinical_indices,
                gex_indices,
                cnv_indices,
                meth_indices,
                mirna_indices,
                mut_indices,
                rppa_indices,
            ]
            # Make sure that all variables are considered in the blocks
            assert sum([len(i) for i in blocks]) == X.shape[1]

            benchmark = {
                "model": [model for i in range(train_splits.shape[0])],
                "concordance": [],
                "ibs": [],
            }
            train_predictions = []
            test_predictions = []
            for i in range(train_splits.shape[0]):
                print(f"Split: {i+1} / 10")
                train_ix = train_splits.iloc[i, :].dropna().values
                test_ix = test_splits.iloc[i, :].dropna().values
                net = HierarchicalSAENet(
                    module=HierarchicalSAE,
                    criterion=hierarchical_sae_criterion,
                    max_epochs=100,
                    lr=0.01,
                    train_split=StratifiedSkorchSurvivalSplit(
                        10, stratified=True
                    ),
                    optimizer=torch.optim.Adam,
                    callbacks=[
                        ("seed", FixRandomSeed(config["seed"])),
                        (
                            "es",
                            EarlyStopping(
                                patience=10,
                                monitor="valid_loss",
                                load_best=True,
                            ),
                        ),
                        (
                            "sched",
                            LRScheduler(
                                torch.optim.lr_scheduler.ReduceLROnPlateau,
                                patience=3,
                                cooldown=5,
                                monitor="valid_loss",
                                verbose=False,
                            ),
                        ),
                    ],
                    verbose=0,
                    batch_size=-1,
                    module__blocks=blocks[:modalities],
                )
                pipe = make_pipeline(StandardScaler(), net)

                pipe.fit(
                    X.iloc[train_ix, :].to_numpy().astype(np.float32),
                    y_str.iloc[train_ix].to_numpy().astype(str),
                )
                train_prediction = pipe.predict(
                    X.iloc[train_ix, :].to_numpy().astype(np.float32),
                )
                test_prediction = pipe.predict(
                    X.iloc[test_ix, :].to_numpy().astype(np.float32),
                )
                train_predictions.append(np.ravel(train_prediction))
                test_predictions.append(np.ravel(test_prediction))
                benchmark["concordance"].append(
                    pipe.score(
                        X.iloc[test_ix, :].to_numpy().astype(np.float32),
                        y_str.iloc[test_ix].to_numpy().astype(str),
                    )
                )
                benchmark["ibs"].append(np.nan)
            pd.DataFrame(benchmark).to_csv(
                f"./data/benchmarks/{cancer}/hierarchical_sae_scores_{modal_mapping[modalities]}_es.csv",
                index=False,
            )
            pd.DataFrame(train_predictions).to_csv(
                f"./data/benchmarks/{cancer}/hierarchical_sae_tcga_train_predictions.csv",
                index=False,
            )
            pd.DataFrame(test_predictions).to_csv(
                f"./data/benchmarks/{cancer}/hierarchical_sae_tcga_test_predictions.csv",
                index=False,
            )


if __name__ == "__main__":
    sys.exit(main())
```
