## Supplementary material for "Clinically driven knowledge distillation for sparsifying high-dimensional multi-omics survival models": Code: run_table_3_students.R

```
library(here)
library(rjson)
library(readr)
library(vroom)
library(dplyr)

source(here::here("src", "utils", "utils.R"))
source(here::here("src", "model", "statistical_models_multi_omics.R"))
config <- rjson::fromJSON(
  file = here::here("config", "config.json")
)
print(config$train_split_name_tcga)

def_order <- c(
  "clinical",
  "gex",
  "mut",
  "mirna",
  "meth",
  "cnv",
  "rppa"
)
modal_mapping <- list(
  "7" = "clinical_mirna",
  "6" = "clinical_cnv",
  "5" = "clinical_rppa",
  "4" = "clinical_meth",
  "3" = "clinical_mut",
  "2" = "clinical_gex",
  "1" = "clinical_only"
)

helper_mapping <- list(
  "7" = c("clinical", "mirna"),
  "6" = c("clinical", "cnv"),
  "5" = c("clinical", "rppa"),
  "4" = c("clinical", "meth"),
  "3" = c("clinical", "mut"),
  "2" = c("clinical", "gex"),
  "1" = c("clinical")
)


for (cancer in config$cancers) {
  print(paste0("Starting cancer: ", cancer))
  set.seed(config$seed)
  data <- data.frame(vroom::vroom(
    here::here(
      "data", "processed", cancer, "merged", config$data_name_tcga
    ),
  ), check.names = FALSE)
  train_splits <- read_csv(
    here::here(
      "data", "splits", cancer, config$train_split_name_tcga
    )
  )

  test_splits <- read_csv(
    here::here(
      "data", "splits", cancer, config$test_split_name_tcga
    )
  )

  train_splits <- format_splits(train_splits)
  test_splits <- format_splits(test_splits)
  results_rb_favoring <- list()
  results_blockforest <- list()
  results_hierarchical_sae <- list()
  results_ensembled_sparse <- list()
  results_ensembled <- list()
  for (modalities in c(2:7)) {
    for (i in 1:length(train_splits)) {
      print(paste0("Split: ", i))
      train_ix <- train_splits[[i]]
      test_ix <- test_splits[[i]]
      X_train <- data[
        train_ix,
        unname(unlist(sapply(helper_mapping[[as.character(modalities)]], function(block) grep(block, colnames(data)))))
      ] %>%
        data.matrix()
      X_test <- data[
        test_ix,
        unname(
          unlist(sapply(helper_mapping[[as.character(modalities)]], function(block) grep(block, colnames(data))))
        )
      ] %>%
        data.matrix()
      y_train <- data.matrix(data[train_ix, 2:1])
      y_test <- data.matrix(data[test_ix, 2:1])
      rb_favoring_train_predictions <- readRDS(here::here("data", "benchmarks", cancer, "randomblock_favoring_10_stratified_all.rds"))[[i]]$train_predictions
      rb_favoring_test_predictions <- readRDS(here::here("data", "benchmarks", cancer, "randomblock_favoring_10_stratified_all.rds"))[[i]]$test_predictions
      blockforest_train_predictions <- readRDS(here::here("data", "benchmarks", cancer, "blockforest_10_stratified_all.rds"))[[i]]$train_predictions
      blockforest_test_predictions <- readRDS(here::here("data", "benchmarks", cancer, "blockforest_10_stratified_all.rds"))[[i]]$test_predictions

      rb_favoring_train_predictions <- rb_favoring_train_predictions[!is.na(rb_favoring_train_predictions)]
      rb_favoring_test_predictions <- rb_favoring_test_predictions[!is.na(rb_favoring_test_predictions)]
      blockforest_train_predictions <- blockforest_train_predictions[!is.na(blockforest_train_predictions)]
      blockforest_test_predictions <- blockforest_test_predictions[!is.na(blockforest_test_predictions)]

      ensembled_train_predictions <- apply(matrix(c(rb_favoring_train_predictions, blockforest_train_predictions), nrow = 2, byrow = TRUE), 2, mean)
      ensembled_test_predictions <- apply(matrix(c(rb_favoring_test_predictions, blockforest_test_predictions), nrow = 2, byrow = TRUE), 2, mean)

      hierarchical_sae_train_predictions <- as.numeric((vroom::vroom(
        here::here(
          "data", "benchmarks", cancer, "hierarchical_sae_tcga_train_predictions.csv"
        )
      ) %>% data.frame())[i, ])
      hierarchical_sae_train_predictions <- hierarchical_sae_train_predictions[!is.na(hierarchical_sae_train_predictions)]
      hierarchical_sae_test_predictions <- as.numeric((vroom::vroom(
        here::here(
          "data", "benchmarks", cancer, "hierarchical_sae_tcga_test_predictions.csv"
        )
      ) %>% data.frame())[i, ])
      hierarchical_sae_test_predictions <- hierarchical_sae_test_predictions[!is.na(hierarchical_sae_test_predictions)]

      eval_points <- sort(unique(y_train[, 1]))
      eval_points <- eval_points[eval_points >= min(y_test[, 1])]
      eval_points <- eval_points[eval_points < max(y_test[, 1])]

      results_rb_favoring[[i]] <- run_kd(
        rb_favoring_train_predictions,
        rb_favoring_test_predictions,
        X_train,
        X_test,
        y_train,
        y_test,
        "BF",
        "clinical" %in% helper_mapping[[as.character(modalities)]] & modalities != 1,
        "lambda.min",
        TRUE
      )

      results_blockforest[[i]] <- run_kd(
        blockforest_train_predictions,
        blockforest_test_predictions,
        X_train,
        X_test,
        y_train,
        y_test,
        "BF",
        "clinical" %in% helper_mapping[[as.character(modalities)]] & modalities != 1,
        "lambda.min",
        TRUE
      )

      results_hierarchical_sae[[i]] <- run_kd(
        hierarchical_sae_train_predictions,
        hierarchical_sae_test_predictions,
        X_train,
        X_test,
        y_train,
        y_test,
        "hierarchical_sae",
        "clinical" %in% helper_mapping[[as.character(modalities)]] & modalities != 1,
        "lambda.min",
        TRUE
      )
    }
    saveRDS(
      results_rb_favoring,
      file = here::here("data", "benchmarks", cancer, paste0("randomblock_favoring_distilled_with_intercept_", modal_mapping[[as.character(modalities)]], ".rds"))
    )
    saveRDS(
      results_blockforest,
      file = here::here("data", "benchmarks", cancer, paste0("blockforest_distilled_with_intercept_", modal_mapping[[as.character(modalities)]], ".rds"))
    )
    saveRDS(
      results_hierarchical_sae,
      file = here::here("data", "benchmarks", cancer, paste0("hierarchical_sae_distilled_with_intercept_", modal_mapping[[as.character(modalities)]], ".rds"))
    )
  }
}
```
