## Supplementary material for "Clinically driven knowledge distillation for sparsifying high-dimensional multi-omics survival models": Code: run_table_4_experiment.R

```
library(here)
library(rjson)
library(readr)
library(vroom)
library(dplyr)

source(here::here("src", "utils", "utils.R"))
source(here::here("src", "model", "statistical_models_multi_omics.R"))
config <- rjson::fromJSON(
  file = here::here("config", "config.json")
)
modal_mapping <- list(
  "7" = "all"
)
naming <- list(
  "1" = "hierarchical_sae_feature_selected",
  "2" = "glmnet_favoring_feature_selected",
  "3" = "rb_favoring_feature_selected",
  "4" = "blockforest_feature_selected",
  "5" = "prioritylasso_feature_selected"
)

for (cancer in config$cancers) {
  print(paste0("Starting cancer: ", cancer))
  set.seed(config$seed)
  data <- data.frame(vroom::vroom(
    here::here(
      "data", "processed", cancer, "merged", config$data_name_tcga
    ),
  ), check.names = FALSE)
  train_splits <- read_csv(
    here::here(
      "data", "splits", cancer, config$train_split_name_tcga
    )
  )
  test_splits <- read_csv(
    here::here(
      "data", "splits", cancer, config$test_split_name_tcga
    )
  )
  hierarchical_sae_coefs <- sapply(readRDS(here::here("data", "benchmarks", cancer, "hierarchical_sae_distilled_all.rds")), function(x) x[[5]])
  rb_favoring_coefs <- sapply(readRDS(here::here("data", "benchmarks", cancer, "randomblock_favoring_distilled_fidelity_with_intercept_all.rds")), function(x) x[[5]])
  blockforest_coefs <- sapply(readRDS(here::here("data", "benchmarks", cancer, "hierarchical_sae_distilled_fidelity_with_intercept_all.rds")), function(x) x[[5]])

  prioritylasso_coefs <- sapply(
    readRDS(here::here("data", "benchmarks", cancer, "prioritylasso_10_stratified_all.rds")),
    function(x) x$selected_variables
  )
  lasso_coefs <- sapply(
    readRDS(here::here("data", "benchmarks", cancer, "glmnet_favor_10_stratified_all.rds")),
    function(x) x$selected_variables
  )

  train_splits <- format_splits(train_splits)
  test_splits <- format_splits(test_splits)
  for (model in c("ridge")) {
    for (modalities in c(7)) {
      results <- list(
        "1" = list(), "2" = list(), "3" = list(), "4" = list(), "5" = list()
      )
      for (z in 1:5) {
        for (i in 1:length(train_splits)) {
          penalty_factor <- NA
          train_ix <- train_splits[[i]]
          test_ix <- test_splits[[i]]
          # For each model and each split,
          # select out the corresponding features
          # and then fit a Ridge model that uses those.
          if (z == 1) {
            X_train <- data[
              train_ix,
              which(colnames(data) %in% hierarchical_sae_coefs[[i]])
            ] %>%
              data.matrix()
            X_test <- data[
              test_ix,
              which(colnames(data) %in% hierarchical_sae_coefs[[i]])
            ] %>%
              data.matrix()
          } else if (z == 2) {
            X_train <- data[
              train_ix,
              which(colnames(data) %in% lasso_coefs[[i]])
            ] %>%
              data.matrix()
            X_test <- data[
              test_ix,
              which(colnames(data) %in% lasso_coefs[[i]])
            ] %>%
              data.matrix()
          } else if (z == 3) {
            X_train <- data[
              train_ix,
              which(colnames(data) %in% rb_favoring_coefs[[i]])
            ] %>%
              data.matrix()
            X_test <- data[
              test_ix,
              which(colnames(data) %in% rb_favoring_coefs[[i]])
            ] %>%
              data.matrix()
          } else if (z == 4) {
            X_train <- data[
              train_ix,
              which(colnames(data) %in% blockforest_coefs[[i]])
            ] %>%
              data.matrix()
            X_test <- data[
              test_ix,
              which(colnames(data) %in% blockforest_coefs[[i]])
            ] %>%
              data.matrix()
          } else if (z == 5) {
            X_train <- data[
              train_ix,
              which(colnames(data) %in% prioritylasso_coefs[[i]])
            ] %>%
              data.matrix()
            colnames(X_train) <- prioritylasso_coefs[[i]]
            X_test <- data[
              test_ix,
              which(colnames(data) %in% prioritylasso_coefs[[i]])
            ] %>%
              data.matrix()
          }
          # If no features were selected by one of the sparse models
          # (singular fit), we assign NA and encode this as random
          # performance (0.5 concordance) and full sparsity
          # when preparing figures and tables.
          if (ncol(X_train) < 1) {
            results[[as.character(z)]][[i]] <- list(
              concordance = NA,
              params = NA,
              selected_variables = NA,
              hazards = NA
            )
            next
          }
          y_train <- data.matrix(data[train_ix, 2:1])
          y_test <- data.matrix(data[test_ix, 2:1])
          def_order <- c(
            "clinical",
            "gex",
            "mut",
            "mirna",
            "meth",
            "cnv",
            "rppa"
          )
          blocks <- list(
            bp1 = grep("clinical", colnames(X_train)),
            bp2 = grep("gex", colnames(X_train)),
            bp3 = grep("mut", colnames(X_train)),
            bp4 = grep("mirna", colnames(X_train)),
            bp5 = grep("meth", colnames(X_train)),
            bp6 = grep("cnv", colnames(X_train)),
            bp7 = grep("rppa", colnames(X_train))
          )

          def_order <- def_order[sapply(blocks, length) > 0]
          blocks <- blocks[sapply(blocks, length) > 0]
          names(blocks) <- paste0(rep("bp", length(blocks)), 1:length(blocks))

          tryCatch(
            {
              print(model)
              results[[as.character(z)]][[i]] <- run_benchmark(
                model = model,
                X_train = X_train,
                y_train = y_train,
                X_test = X_test,
                y_test = y_test,
                blocks,
                seed = config$seed
              )
            },
            error = function(cond) {
              # In the case of failure, we assign NA values
              # which we later replace with random performance
              # for concordance and complete sparsity for
              # the number of features used.
              # Note: Need to make sure to use
              # global assignment here.
              print(cond)
              print("Error running model.")
              print("Saving everything as NA.")
              results[[as.character(z)]][[i]] <<- list(
                concordance = NA,
                params = NA,
                selected_variables = NA,
                hazards = NA
              )
            }
          )
        }
        saveRDS(
          results[[as.character(z)]],
          file = here::here("data", "benchmarks", cancer, paste0(model, paste0("_10_stratified_", modal_mapping[as.character(modalities)], "_", naming[[as.character(z)]], ".rds")))
        )
      }
    }
  }
}
```
