## Supplementary material for "Clinically driven knowledge distillation for sparsifying high-dimensional multi-omics survival models": Code: autoencoders.py

```
#!/usr/bin/env python
# -*- coding: utf-8 -*-


import os
import sys

module_path = os.path.abspath(os.path.join("./src/"))
if module_path not in sys.path:
    sys.path.append(module_path)

import math

import numpy as np
import skorch
import torch
from torch import nn

from model.model_utils import BaseSurvivalNeuralNet

# All code adapted from:
# Wissel, D., Rowson, D., and Boeva, V. Hierarchical autoencoder-based integration improves performance in multi-omics cancer survival models through soft modality selection. bioRxiv, 2022.
class Encoder(nn.Module):
    """Base class modeling the encoder portion of an autoencoder.

    Attributes:
        input_dimension: Input size going into the encoder.
        hidden_layer_size: Number of hidden nodes within each hidden layer of the encoder.
        activation: Non-linear activation method to be used by the encoder.
        hidden_layers: Number of hidden layers within the encoder.
        embedding_dimension: Dimensionality of the final output of the encoder (i.e., the latent space).
        encode: `torch.Sequential` module containing the full encoder.
    """

    def __init__(
        self,
        input_dimension,
        hidden_layer_size=128,
        activation=nn.PReLU,
        hidden_layers=1,
        embedding_dimension=64,
    ):
        super().__init__()
        encoder = []
        current_size = input_dimension
        next_size = hidden_layer_size
        for i in range(hidden_layers):
            if i != 0:
                current_size = next_size
                # Slowly halve size of the AE hidden layer dimension
                # over time until we reach the embedding dimension.
                # Take max since we do not want the non-bottleneck
                # layers to become smaller than the bottleneck.
                next_size = max(int(next_size / 2), embedding_dimension)
            encoder.append(nn.Linear(current_size, next_size))
            encoder.append(activation())
            encoder.append(nn.BatchNorm1d(next_size))
        if hidden_layers > 0:
            encoder.append(nn.Linear(next_size, embedding_dimension))
        else:
            encoder.append(nn.Linear(input_dimension, embedding_dimension))
        # Do not include an activation before the embedding.
        self.encode = nn.Sequential(*encoder)
        self.input_dimension = input_dimension
        self.hidden_layer_size = hidden_layer_size
        self.activation = activation
        self.hidden_layers = hidden_layers
        self.embedding_dimension = embedding_dimension

    def forward(self, x):
        return self.encode(x)


class Decoder(nn.Module):
    """Base class modeling the decoder portion of an autoencoder.

    Attributes:
        decode: `torch.Sequential` module containing the full decoder.
    """

    def __init__(self, encoder, activation=nn.PReLU):
        super().__init__()
        decoder = []
        # Build up the decoder symmetrically from the encoder.
        for layer in encoder.encode[::-1][::3]:
            current_size = layer.weight.shape[0]
            next_size = layer.weight.shape[1]
            decoder.append(nn.BatchNorm1d(current_size))
            decoder.append(nn.Linear(current_size, next_size))
            decoder.append(activation())

        # Remove the embedding before the output.
        self.decode = nn.Sequential(*(decoder[:-1]))

    def forward(self, x):
        return self.decode(x)


class AE(nn.Module):
    """Base class modeling an autoencoder.

    Attributes:
        encode: `torch.Sequential` module containing the full encoder.
        decode: `torch.Sequential` module containing the full decoder.
        input_dimension: Input size going into the encoder.
        hidden_layer_size: Number of hidden nodes within each hidden layer of the encoder.
        activation: Non-linear activation method to be used by the encoder.
        hidden_layers: Number of hidden layers within the encoder.
        embedding_dimension: Dimensionality of the final output of the encoder (i.e., the latent space).
    """

    def __init__(
        self,
        input_dimension,
        hidden_layer_size=128,
        activation=nn.PReLU,
        hidden_layers=1,
        embedding_dimension=64,
    ):
        super().__init__()
        self.encode = Encoder(
            input_dimension,
            hidden_layer_size,
            activation,
            hidden_layers,
            embedding_dimension,
        )
        self.decode = Decoder(self.encode, activation)
        self.input_dimension = input_dimension
        self.hidden_layer_size = hidden_layer_size
        self.activation = activation
        self.hidden_layers = hidden_layers
        self.embedding_dimension = embedding_dimension

    def forward(self, x):
        encoded = self.encode(x)
        decoded = self.decode(encoded)
        return encoded, decoded


class HazardRegression(nn.Module):
    """Base class modeling a cox hazard regression problem.

    Attributes:
        input_dimension: Number of covariates to be input to the cox regression.
        hidden_layer_size: Number of hidden nodes within each hidden layer of the regression.
        activation: Non-linear activation to be used after each hidden layer of the regression.
        hidden_layers: Number of hidden layers to be used within the regression.
    """

    def __init__(
        self,
        input_dimension,
        hidden_layer_size=32,
        activation=nn.PReLU,
        hidden_layers=0,
    ):
        super().__init__()
        hazard = []
        current_size = input_dimension
        for layer in range(hidden_layers):
            next_size = hidden_layer_size
            hazard.append(nn.BatchNorm1d(current_size))
            hazard.append(nn.Linear(current_size, next_size))
            hazard.append(activation())
            current_size = next_size
        # Batch norm the embedding before prediction.
        hazard.append(nn.BatchNorm1d(current_size))
        hazard.append(nn.Linear(current_size, 1))
        self.hazard = nn.Sequential(*hazard)

    def forward(self, x):
        return self.hazard(x)


class HierarchicalSAE(nn.Module):
    """Implements a hierarchical supervised multi-modal autoencoder."""

    def __init__(
        self,
        blocks,
        block_embedding_dimension=64,
        block_hidden_layers=1,
        block_hidden_layer_size=128,
        common_embedding_dimension=64,
        common_hidden_layers=0,
        common_hidden_layer_size=128,
        block_activation=nn.PReLU,
        common_activation=nn.PReLU,
        hazard_hidden_layer_size=64,
        hazard_activation=nn.PReLU,
        hazard_hidden_layers=0,
        lambda_q=0.001,
        supervise_aes=True,
    ):
        super().__init__()
        block_aes = []
        block_hazards = []

        for ix, block in enumerate(blocks):
            block_aes.append(
                AE(
                    len(block),
                    block_hidden_layer_size,
                    block_activation,
                    block_hidden_layers,
                    block_embedding_dimension,
                )
            )
            block_hazards.append(
                HazardRegression(
                    block_embedding_dimension,
                    hazard_hidden_layer_size,
                    hazard_activation,
                    hazard_hidden_layers,
                )
            )
        self.block_hazards = nn.ModuleList(block_hazards)
        self.block_aes = nn.ModuleList(block_aes)
        self.ae = AE(
            block_embedding_dimension * len(blocks),
            common_hidden_layer_size,
            common_activation,
            common_hidden_layers,
            common_embedding_dimension,
        )
        self.hazard = HazardRegression(
            common_embedding_dimension,
            hazard_hidden_layer_size,
            hazard_activation,
            hazard_hidden_layers,
        )
        self.blocks = blocks
        self.block_embedding_dimension = block_embedding_dimension
        self.block_hidden_layers = block_hidden_layers
        self.block_hidden_layer_size = block_hidden_layer_size
        self.common_embedding_dimension = common_embedding_dimension
        self.common_hidden_layers = common_hidden_layers
        self.common_hidden_layer_size = common_hidden_layer_size
        self.block_activation = block_activation
        self.common_activation = common_activation
        self.lambda_q = lambda_q
        self.supervise_aes = supervise_aes

    def forward(self, x):
        # Separate initial blocks, also needed for calculating
        # the reconstruction loss later.
        original_blocks = [x[:, block] for block in self.blocks]
        # Run first level autoencoders.
        block_aes = [
            self.block_aes[ix](x[:, block])
            for ix, block in enumerate(self.blocks)
        ]
        # Get first level autoencoder predicted partial hazards
        # in order to calculate the cox losses for each
        # first level supervivsed autoencoder.
        block_hazards = [
            self.block_hazards[ix](i[0]) for ix, i in enumerate(block_aes)
        ]
        # Get first level decoded representations.
        blocks_decoded = [i[1] for i in block_aes]

        # Concatenate first level latent spaces in order to put
        # them into the second level autoencoder.
        original_common = torch.cat([i[0] for i in block_aes], dim=1)
        ae = self.ae(original_common)
        decoded = ae[1]
        hazard = self.hazard(ae[0])

        # Return various things needed for the loss function.
        return (
            hazard,
            original_common,
            decoded,
            original_blocks,
            blocks_decoded,
            block_hazards,
            ae[0],
        )


class HierarchicalSAENet(BaseSurvivalNeuralNet):
    def get_loss(self, y_pred, y_true, X=None, training=False):
        losses = self.criterion_(
            y_pred,
            y_true,
            torch.tensor(np.ones(len(y_true))),
            len(self.module_.blocks),
        )
        # Calculate weight decay only on linear layers.
        weight_decay = 0
        for i in self.module_.block_aes:
            for q in [z for z in i.encode.encode if isinstance(z, nn.Linear)]:
                weight_decay += torch.square(torch.norm(q.weight, p=2))
            for q in [z for z in i.decode.decode if isinstance(z, nn.Linear)]:
                weight_decay += torch.square(torch.norm(q.weight, p=2))

        for i in [
            q
            for q in self.module_.ae.encode.encode
            if isinstance(q, nn.Linear)
        ]:
            weight_decay += torch.square(torch.norm(i.weight, p=2))
        for i in [
            q
            for q in self.module_.ae.decode.decode
            if isinstance(q, nn.Linear)
        ]:
            weight_decay += torch.square(torch.norm(i.weight, p=2))
        for i in [
            q for q in self.module_.hazard.hazard if isinstance(q, nn.Linear)
        ]:
            weight_decay += torch.square(
                torch.norm(
                    i.weight,
                    p=2,
                )
            )
        total_reconstruction_loss = 0
        total_block_hazards = 0
        # Sum up total reconstruction loss as well as block partial
        # hazard losses.
        for key, val in losses.items():
            if key != "cox" and "hazard" not in key:
                total_reconstruction_loss += val

        for key, val in losses.items():
            if "hazard" in key:
                total_block_hazards += val
        if self.module_.supervise_aes:
            return (
                losses["cox"]
                + total_reconstruction_loss
                + total_block_hazards
                + self.module_.lambda_q * weight_decay
            )
        else:
            return (
                losses["cox"]
                + total_reconstruction_loss
                + self.module_.lambda_q * weight_decay
            )


class ConcatSAE(nn.Module):
    def __init__(
        self,
        blocks,
        block_embedding_dimension=64,
        block_hidden_layers=1,
        block_hidden_layer_size=128,
        block_activation=nn.PReLU,
        hazard_hidden_layer_size=64,
        hazard_activation=nn.PReLU,
        hazard_hidden_layers=0,
        lambda_q=0.001,
        supervise_aes=True,
    ):
        super().__init__()
        block_aes = []
        block_hazards = []

        for block in blocks:
            block_aes.append(
                AE(
                    len(block),
                    block_hidden_layer_size,
                    block_activation,
                    block_hidden_layers,
                    block_embedding_dimension,
                )
            )
            block_hazards.append(
                HazardRegression(
                    block_embedding_dimension,
                    hazard_hidden_layer_size,
                    hazard_activation,
                    hazard_hidden_layers,
                )
            )
        self.block_hazards = nn.ModuleList(block_hazards)
        self.block_aes = nn.ModuleList(block_aes)
        self.hazard = HazardRegression(
            block_embedding_dimension * len(blocks),
            hazard_hidden_layer_size,
            hazard_activation,
            hazard_hidden_layers,
        )
        self.blocks = blocks
        self.block_embedding_dimension = block_embedding_dimension
        self.block_hidden_layers = block_hidden_layers
        self.block_hidden_layer_size = block_hidden_layer_size
        self.lambda_q = lambda_q
        self.supervise_aes = supervise_aes

    def forward(self, x):
        original_blocks = [x[:, block] for block in self.blocks]
        block_aes = [
            self.block_aes[ix](x[:, block])
            for ix, block in enumerate(self.blocks)
        ]
        original_common = torch.cat([i[0] for i in block_aes], dim=1)
        block_hazards = [
            self.block_hazards[ix](i[0]) for ix, i in enumerate(block_aes)
        ]
        blocks_decoded = [i[1] for i in block_aes]
        hazard = self.hazard(original_common)
        return (
            hazard,
            original_blocks,
            blocks_decoded,
            block_hazards,
            original_common,
        )


class ConcatSAENet(BaseSurvivalNeuralNet):
    def get_loss(self, y_pred, y_true, X=None, training=False):
        losses = self.criterion_(
            y_pred,
            y_true,
            torch.tensor(np.ones(len(y_true))),
            len(self.module_.blocks),
        )
        weight_decay = 0
        for i in self.module_.block_aes:
            for q in [z for z in i.encode.encode if isinstance(z, nn.Linear)]:
                weight_decay += torch.square(torch.norm(q.weight, p=2))
            for q in [z for z in i.decode.decode if isinstance(z, nn.Linear)]:
                weight_decay += torch.square(torch.norm(q.weight, p=2))
        for i in [
            q for q in self.module_.hazard.hazard if isinstance(q, nn.Linear)
        ]:
            weight_decay += torch.square(
                torch.norm(
                    i.weight,
                    p=2,
                )
            )
        total_reconstruction_loss = 0
        for key, val in losses.items():
            if key != "cox" and "hazard" not in key:
                total_reconstruction_loss += val
        total_block_hazards = 0
        for key, val in losses.items():
            if "hazard" in key:
                total_block_hazards += val
        if self.module_.supervise_aes:
            return (
                losses["cox"]
                + total_reconstruction_loss
                + total_block_hazards
                + self.module_.lambda_q * weight_decay
            )
        else:
            return (
                losses["cox"]
                + total_reconstruction_loss
                + self.module_.lambda_q * weight_decay
            )


class PoolSAE(nn.Module):
    def __init__(
        self,
        blocks,
        beta=1,
        block_embedding_dimension=64,
        block_hidden_layers=1,
        block_hidden_layer_size=128,
        block_activation=nn.PReLU,
        hazard_hidden_layer_size=64,
        hazard_activation=nn.PReLU,
        hazard_hidden_layers=0,
        lambda_q=0.001,
        fusion="mean",
        supervise_aes=True,
    ):
        super().__init__()
        block_aes = []
        block_hazards = []

        for block in blocks:
            block_aes.append(
                AE(
                    len(block),
                    block_hidden_layer_size,
                    block_activation,
                    block_hidden_layers,
                    block_embedding_dimension,
                )
            )
            block_hazards.append(
                HazardRegression(
                    block_embedding_dimension,
                    hazard_hidden_layer_size,
                    hazard_activation,
                    hazard_hidden_layers,
                )
            )
        self.block_hazards = nn.ModuleList(block_hazards)
        self.block_aes = nn.ModuleList(block_aes)
        self.hazard = HazardRegression(
            block_embedding_dimension,
            hazard_hidden_layer_size,
            hazard_activation,
            hazard_hidden_layers,
        )
        self.blocks = blocks
        self.beta = beta
        self.block_embedding_dimension = block_embedding_dimension
        self.block_hidden_layers = block_hidden_layers
        self.block_hidden_layer_size = block_hidden_layer_size
        self.block_activation = block_activation
        self.fusion = fusion
        self.lambda_q = lambda_q
        self.supervise_aes = supervise_aes

    def forward(self, x):
        original_blocks = [x[:, block] for block in self.blocks]
        block_aes = [
            self.block_aes[ix](x[:, block])
            for ix, block in enumerate(self.blocks)
        ]
        if self.fusion == "mean":
            original_common = torch.mean(
                torch.stack([i[0] for i in block_aes]), dim=0
            )
        else:
            original_common = torch.max(
                torch.stack([i[0] for i in block_aes]), dim=0
            )[0]
        block_hazards = [
            self.block_hazards[ix](i[0]) for ix, i in enumerate(block_aes)
        ]
        blocks_decoded = [i[1] for i in block_aes]
        hazard = self.hazard(original_common)
        return (
            hazard,
            original_blocks,
            blocks_decoded,
            block_hazards,
            original_common,
        )


class MeanSAEEncodeOnly(nn.Module):
    def __init__(
        self,
        blocks,
        beta=1,
        block_embedding_dimension=64,
        block_hidden_layers=1,
        block_hidden_layer_size=128,
        common_embedding_dimension=64,
        common_hidden_layers=0,
        common_hidden_layer_size=128,
        block_activation=nn.PReLU,
        common_activation=nn.PReLU,
        hazard_hidden_layer_size=64,
        hazard_activation=nn.PReLU,
        hazard_hidden_layers=0,
        lambda_q=0.001,
        fusion="mean",
    ):
        super().__init__()
        block_aes = []
        block_hazards = []

        for block in blocks:
            block_aes.append(
                AE(
                    len(block),
                    block_hidden_layer_size,
                    block_activation,
                    block_hidden_layers,
                    block_embedding_dimension,
                )
            )
            block_hazards.append(
                HazardRegression(
                    block_embedding_dimension,
                    hazard_hidden_layer_size,
                    hazard_activation,
                    hazard_hidden_layers,
                )
            )
        self.block_hazards = nn.ModuleList(block_hazards)
        self.block_aes = nn.ModuleList(block_aes)
        self.encoder = Encoder(
            block_embedding_dimension,
            common_hidden_layer_size,
            common_activation,
            common_hidden_layers,
            common_embedding_dimension,
        )
        self.hazard = HazardRegression(
            block_embedding_dimension,
            hazard_hidden_layer_size,
            hazard_activation,
            hazard_hidden_layers,
        )
        self.blocks = blocks
        self.beta = beta
        self.block_embedding_dimension = block_embedding_dimension
        self.block_hidden_layers = block_hidden_layers
        self.block_hidden_layer_size = block_hidden_layer_size
        self.common_embedding_dimension = common_embedding_dimension
        self.common_hidden_layers = common_hidden_layers
        self.common_hidden_layer_size = common_hidden_layer_size
        self.block_activation = block_activation
        self.common_activation = common_activation
        self.lambda_ = lambda_q
        self.fusion = fusion

    def forward(self, x):
        original_blocks = [x[:, block] for block in self.blocks]
        block_aes = [
            self.block_aes[ix](x[:, block])
            for ix, block in enumerate(self.blocks)
        ]
        if self.fusion == "mean":
            original_common = torch.mean(
                torch.stack([i[0] for i in block_aes]), dim=0
            )
        else:
            original_common = torch.max(
                torch.stack([i[0] for i in block_aes]), dim=0
            )[0]
        block_hazards = [
            self.block_hazards[ix](i[0]) for ix, i in enumerate(block_aes)
        ]
        blocks_decoded = [i[1] for i in block_aes]
        hazard = self.hazard(self.encoder(original_common))
        return (
            hazard,
            original_blocks,
            blocks_decoded,
            block_hazards,
            original_common,
        )


class MeanSAENet(BaseSurvivalNeuralNet):
    def get_loss(self, y_pred, y_true, X=None, training=False):
        losses = self.criterion_(
            y_pred,
            y_true,
            torch.tensor(np.ones(len(y_true))),
            len(self.module_.blocks),
        )
        weight_decay = 0
        for i in self.module_.block_aes:
            for q in [z for z in i.encode.encode if isinstance(z, nn.Linear)]:
                weight_decay += torch.square(torch.norm(q.weight, p=2))
            for q in [z for z in i.decode.decode if isinstance(z, nn.Linear)]:
                weight_decay += torch.square(torch.norm(q.weight, p=2))
        for q in [
            z for z in self.module_.hazard.hazard if isinstance(z, nn.Linear)
        ]:
            weight_decay += torch.square(torch.norm(q.weight, p=2))

        total_reconstruction_loss = 0
        for key, val in losses.items():
            if key != "cox" and "hazard" not in key:
                total_reconstruction_loss += val
        total_block_hazards = 0
        for key, val in losses.items():
            if "hazard" in key:
                total_block_hazards += val
        if self.module_.supervise_aes:
            return (
                losses["cox"]
                + total_reconstruction_loss
                + total_block_hazards
                + self.module_.lambda_q * weight_decay
            )
        else:
            return (
                losses["cox"]
                + total_reconstruction_loss
                + self.module_.lambda_q * weight_decay
            )


class MSAEEncodeOnlyNet(BaseSurvivalNeuralNet):
    def get_loss(self, y_pred, y_true, X=None, training=False):
        losses = self.criterion_(
            y_pred,
            y_true,
            torch.tensor(np.ones(len(y_true))),
            len(self.module_.blocks),
        )
        weight_decay = 0
        for i in self.module_.block_aes:
            for q in [z for z in i.encode.encode if isinstance(z, nn.Linear)]:
                weight_decay += torch.square(torch.norm(q.weight, p=2))
            for q in [z for z in i.decode.decode if isinstance(z, nn.Linear)]:
                weight_decay += torch.square(torch.norm(q.weight, p=2))
        for i in [
            q for q in self.module_.encoder.encode if isinstance(q, nn.Linear)
        ]:
            weight_decay += torch.square(torch.norm(i.weight, p=2))
        for q in [
            z for z in self.module_.hazard.hazard if isinstance(z, nn.Linear)
        ]:
            weight_decay += torch.square(torch.norm(q.weight, p=2))

        total_reconstruction_loss = 0
        for key, val in losses.items():
            if key != "cox" and "hazard" not in key:
                total_reconstruction_loss += val
        total_block_hazards = 0
        for key, val in losses.items():
            if "hazard" in key:
                total_block_hazards += val
        return (
            losses["cox"]
            + total_reconstruction_loss
            + total_block_hazards
            + self.module_.lambda_q * weight_decay
        )


class HierarchicalSAEEncodeOnly(nn.Module):
    def __init__(
        self,
        blocks,
        block_embedding_dimension=64,
        block_hidden_layers=1,
        block_hidden_layer_size=128,
        common_embedding_dimension=64,
        common_hidden_layers=0,
        common_hidden_layer_size=128,
        block_activation=nn.PReLU,
        common_activation=nn.PReLU,
        hazard_hidden_layer_size=64,
        hazard_activation=nn.PReLU,
        hazard_hidden_layers=0,
        lambda_q=0.001,
    ):
        super().__init__()
        block_aes = []
        block_hazards = []
        for ix, block in enumerate(blocks):
            block_aes.append(
                AE(
                    len(block),
                    block_hidden_layer_size,
                    block_activation,
                    block_hidden_layers,
                    block_embedding_dimension,
                )
            )
            block_hazards.append(
                HazardRegression(
                    block_embedding_dimension,
                    hazard_hidden_layer_size,
                    hazard_activation,
                    hazard_hidden_layers,
                )
            )
        self.block_hazards = nn.ModuleList(block_hazards)
        self.block_aes = nn.ModuleList(block_aes)
        self.encoder = Encoder(
            block_embedding_dimension * len(blocks),
            common_hidden_layer_size,
            common_activation,
            common_hidden_layers,
            common_embedding_dimension,
        )
        self.hazard = HazardRegression(
            common_embedding_dimension,
            hazard_hidden_layer_size,
            hazard_activation,
            hazard_hidden_layers,
        )
        self.blocks = blocks
        self.block_embedding_dimension = block_embedding_dimension
        self.block_hidden_layers = block_hidden_layers
        self.block_hidden_layer_size = block_hidden_layer_size
        self.common_embedding_dimension = common_embedding_dimension
        self.common_hidden_layers = common_hidden_layers
        self.common_hidden_layer_size = common_hidden_layer_size
        self.block_activation = block_activation
        self.common_activation = common_activation
        self.lambda_ = lambda_q

    def forward(self, x):
        original_blocks = [x[:, block] for block in self.blocks]
        block_aes = [
            self.block_aes[ix](x[:, block])
            for ix, block in enumerate(self.blocks)
        ]
        original_common = torch.cat([i[0] for i in block_aes], dim=1)
        block_hazards = [
            self.block_hazards[ix](i[0]) for ix, i in enumerate(block_aes)
        ]
        blocks_decoded = [i[1] for i in block_aes]
        encoded = self.encoder(original_common)
        hazard = self.hazard(encoded)

        return (
            hazard,
            original_blocks,
            blocks_decoded,
            block_hazards,
            encoded,
            original_common,
        )


class HierarchicalSAEEncodeOnlyNet(BaseSurvivalNeuralNet):
    def get_loss(self, y_pred, y_true, X=None, training=False):
        losses = self.criterion_(
            y_pred,
            y_true,
            torch.tensor(np.ones(len(y_true))),
            len(self.module_.blocks),
        )
        weight_decay = 0
        for i in self.module_.block_aes:
            for q in [z for z in i.encode.encode if isinstance(z, nn.Linear)]:
                weight_decay += torch.square(torch.norm(q.weight, p=2))
            for q in [z for z in i.decode.decode if isinstance(z, nn.Linear)]:
                weight_decay += torch.square(torch.norm(q.weight, p=2))

        for i in [
            q for q in self.module_.encoder.encode if isinstance(q, nn.Linear)
        ]:
            weight_decay += torch.square(torch.norm(i.weight, p=2))
        for i in [
            q for q in self.module_.hazard.hazard if isinstance(q, nn.Linear)
        ]:
            weight_decay += torch.square(
                torch.norm(
                    i.weight,
                    p=2,
                )
            )
        total_reconstruction_loss = 0
        for key, val in losses.items():
            if key != "cox" and "hazard" not in key:
                total_reconstruction_loss += val
        total_block_hazards = 0
        for key, val in losses.items():
            if "hazard" in key:
                total_block_hazards += val
        return (
            losses["cox"]
            + total_reconstruction_loss
            + total_block_hazards
            + self.module_.lambda_q * weight_decay
        )
```
