## Supplementary material for "Clinically driven knowledge distillation for sparsifying high-dimensional multi-omics survival models": Code: model_utils.py

```
#!/usr/bin/env python
# -*- coding: utf-8 -*-

import numpy as np
from lifelines.utils import concordance_index
from skorch import NeuralNet


# All code adapted from:
# Wissel, D., Rowson, D., and Boeva, V. Hierarchical autoencoder-based integration improves performance in multi-omics cancer survival models through soft modality selection. bioRxiv, 2022.
class BaseSurvivalNeuralNet(NeuralNet):
    def score(self, X: np.ndarray, y: np.ndarray):
        try:
            concordance = concordance_index(
                event_times=np.vstack(np.char.split(np.array(y), sep="|"))[
                    :, 1
                ].astype(np.float32),
                # Flip the scoring since `lifelines` calculates
                # the concordance slightly differently, refer to
                # https://stats.stackexchange.com/questions/352183/use-median-survival-time-to-calculate-cph-c-statistic/352435#352435
                predicted_scores=np.negative(np.squeeze(self.predict(X))),
                event_observed=np.vstack(np.char.split(np.array(y), sep="|"))[
                    :, 0
                ].astype(int),
            )
        except ValueError:
            # In case of error, return 0.5
            # concordance (= random guess).
            concordance = 0.5
        return concordance
```
