## Supplementary material for "Clinically driven knowledge distillation for sparsifying high-dimensional multi-omics survival models": Code: statistical_models_multi_omics.R

```
library(mlr3proba)
library(here)
library(blockForest)
library(glmnet)
library(coefplot)
library(ranger)
library(tuneRanger)
library(tictoc)
library(readr)
library(vroom)
library(Hmisc)
library(prioritylasso)
library(prodlim)
library(ipred)
library(pec)

run_kd <- function(train_predictions, test_predictions, X_train, X_test, y_train, y_test, teacher, fit_unpenalized_clinical, s = "lambda.min", fit_intercept = FALSE) {
  set.seed(42)
  # Prepare penalty factors to fit model with unpenalized clinical
  # variables.
  pf <- rep(1, ncol(X_train))
  pf[grep("clinical", colnames(X_train))] <- 0
  nFolds <- 5
  foldid <- sample(rep(seq(nFolds), length.out = nrow(X_train)))
  if (fit_unpenalized_clinical) {
    learner_kd <- cv.glmnet(
      x = X_train,
      y = train_predictions,
      nfolds = nFolds,
      foldid = foldid,
      alpha = 1,
      penalty.factor = pf,
      intercept = fit_intercept
    )
    kd_favor_clinical_train_predictions <- predict(learner_kd,
      newx = X_train,
      s = s, arg = "response"
    )[, 1]
    kd_favor_clinical_test_predictions <- predict(learner_kd,
      newx = X_test,
      s = s, arg = "response"
    )[, 1]
    kd_favor_clinical_concordance <- unname(rcorr.cens(
      -kd_favor_clinical_test_predictions,
      Surv(y_test)
    )[1])
    kd_favor_clinical_selected_variables <- rownames(extract.coef(learner_kd, lambda = s))
  } else {
    kd_favor_clinical_concordance <- NA
    kd_favor_clinical_selected_variables <- NA
  }
  pf <- rep(1, ncol(X_train))
  learner_kd <- cv.glmnet(
    x = X_train,
    y = train_predictions,
    nfolds = nFolds,
    foldid = foldid,
    alpha = 1,
    penalty.factor = pf,
    intercept = fit_intercept
  )
  kd_train_predictions <- predict(learner_kd,
    newx = X_train,
    s = s, arg = "response"
  )[, 1]
  kd_test_predictions <- predict(learner_kd,
    newx = X_test,
    s = s, arg = "response"
  )[, 1]
  kd_concordance <- unname(rcorr.cens(
    -kd_test_predictions,
    Surv(y_test)
  )[1])
  kd_selected_variables <- rownames(extract.coef(learner_kd, lambda = s))

  # Calculate student-teacher fidelity.
  teacher_student_train_rank_correlation <- cor(
    x = train_predictions,
    y = kd_train_predictions,
    method = "spearman"
  )

  teacher_student_test_rank_correlation <- cor(
    x = test_predictions,
    y = kd_test_predictions,
    method = "spearman"
  )
  if (fit_unpenalized_clinical) {
    teacher_favor_student_train_rank_correlation <- cor(
      x = train_predictions,
      y = kd_favor_clinical_train_predictions,
      method = "spearman"
    )

    teacher_favor_student_test_rank_correlation <- cor(
      x = test_predictions,
      y = kd_favor_clinical_test_predictions,
      method = "spearman"
    )
  }


  if (fit_unpenalized_clinical) {
    list(
      kd_concordance,
      kd_selected_variables,
      list(
        teacher_student_train_rank_correlation,
        teacher_student_test_rank_correlation
      ),
      kd_favor_clinical_concordance,
      kd_favor_clinical_selected_variables,
      list(
        teacher_favor_student_train_rank_correlation,
        teacher_favor_student_test_rank_correlation
      )
    )
  } else {
    # If no clinically guided distillation was fit, we
    # only return the results for the standard student.
    list(
      kd_concordance,
      kd_selected_variables,
      list(
        teacher_student_train_rank_correlation,
        teacher_student_test_rank_correlation
      )
    )
  }
}

# Adapted from ipred::ibrier to work with our data structure to calculate
# the integrated brier score.
ibrier <- function(obj, pred, btime = range(obj[, 1])) {
  if (!inherits(obj, "Surv")) {
    stop("obj is not of class Surv")
  }
  class(obj) <- NULL
  if (attr(obj, "type") != "right") {
    stop("only right-censoring allowed")
  }
  N <- nrow(obj)
  time <- obj[, 1]
  ot <- order(time)
  cens <- obj[ot, 2]
  time <- time[ot]
  if (is.null(btime)) {
    stop("btime not given")
  }
  if (length(btime) < 1) {
    stop("btime not given")
  }
  if (length(btime) == 2) {
    if (btime[1] < min(time)) {
      warning("btime[1] is smaller than min(time)")
    }
    if (btime[2] > max(time)) {
      warning("btime[2] is larger than max(time)")
    }
    btime <- time[time >= btime[1] & time <= btime[2]]
  }
  ptype <- class(pred)[1]
  if (is.null(ptype)) {
    if (is.vector(pred)) {
      ptype <- "vector"
    }
    if (is.list(pred)) {
      ptype <- "list"
    }
  }
  if (ptype == "numeric" && is.vector(pred)) {
    ptype <- "vector"
  }
  survs <- NULL
  switch(ptype,
    survfit = {
      survs <- getsurv(pred, btime)
      survs <- matrix(rep(survs, N), nrow = length(btime))
    },
    list = {
      if (!inherits(pred[[1]], "survfit")) stop("pred is not a list of survfit objects")
      if (length(pred) != N) stop("pred must be of length(time)")
      pred <- pred[ot]
      survs <- matrix(unlist(lapply(pred, getsurv, times = btime)),
        nrow = length(btime), ncol = N
      )
    },
    vector = {
      if (length(pred) != N) stop("pred must be of length(time)")
      if (length(btime) != 1) stop("cannot compute integrated Brier score with pred")
      survs <- pred[ot]
    },
    matrix = {
      if (all(dim(pred) == c(length(btime), N))) {
        survs <- pred[
          ,
          ot
        ]
      } else {
        stop("wrong dimensions of pred")
      }
    }
  )
  if (is.null(survs)) {
    stop("unknown type of pred")
  }
  hatcdist <- prodlim(Surv(time, cens) ~ 1, reverse = TRUE)
  csurv <- predict(hatcdist, times = time, type = "surv")
  csurv[csurv == 0] <- Inf
  csurv_btime <- predict(hatcdist, times = btime, type = "surv")
  csurv_btime[is.na(csurv_btime)] <- min(csurv_btime, na.rm = TRUE)
  csurv_btime[csurv_btime == 0] <- Inf
  bsc <- rep(0, length(btime))
  if (length(btime) > 1) {
    for (j in 1:length(btime)) {
      help1 <- as.integer(time <= btime[j] & cens == 1)
      help2 <- as.integer(time > btime[j])
      bsc[j] <- mean((0 - survs[j, ])^2 * help1 * (1 / csurv) +
        (1 - survs[j, ])^2 * help2 * (1 / csurv_btime[j]))
    }
    idx <- 2:length(btime)
    RET <- diff(btime) %*% ((bsc[idx - 1] + bsc[idx]) / 2)
    RET <- RET / diff(range(btime))
    names(RET) <- "integrated Brier score"
    attr(RET, "time") <- range(btime)
  } else {
    help1 <- as.integer(time <= btime & cens == 1)
    help2 <- as.integer(time > btime)
    cs <- predict(hatcdist, times = btime, type = "surv")
    if (cs == 0) {
      cs <- Inf
    }
    RET <- mean((0 - survs)^2 * help1 * (1 / csurv) + (1 -
      survs)^2 * help2 * (1 / cs))
    names(RET) <- "Brier score"
    attr(RET, "time") <- btime
  }
  RET
}


#' Run multi-omics survival benchmark as per the requirements of our study.
#'
#' @description
#' `run_benchmark()` allows for easy benchmarking, as we can abstract
#' all the model specific code handling away to live here and can let
#' our main driver script not worry about subtle API differences between models.
#' @param model String specifiying which model is to be run.
#' @param X_train nxp design training matrix.
#' @param y_train nx1 `Surv` survival training targets.
#' @param X_test nxp design test matrix.
#' @param y_test nx1 `Surv` survival test targets.
#' @param blocks List specifying which block (i.e., modality) each input variable
#' belongs to.
#' @param seed Integer specifying the random seed to be set for reproducibility.
run_benchmark <- function(model,
                          X_train,
                          y_train,
                          X_test,
                          y_test,
                          blocks,
                          seed = 42) {
  allowable_models <- c(
    "blockforest",
    "randomblock_favoring",
    "prioritylasso",
    "ridge",
    "glmnet_favor",
    "clinical_ridge"
  )
  if (!model %in% allowable_models) {
    stop("Model chosen not available. Please check the documentation for allowable models.")
  }
  if (model %in% c("blockforest", "randomblock_favoring")) {
    set.seed(seed)
    block.method <- "BlockForest"
    always.select.block <- 0
    if (model == "randomblock_favoring") {
      block.method <- "RandomBlock"
      # Favor clinical variables for RandomBlock.
      # Thus, have to make sure that clinical is always the first
      # block, but we handle this.
      always.select.block <- 1
    }
    tic()
    learner <- blockfor(
      X = X_train,
      y = y_train,
      blocks = blocks,
      block.method = block.method,
      always.select.block = always.select.block,
      seed = seed,
      verbose = FALSE
    )
    exectime <- toc()
    exectime <- exectime$toc - exectime$tic
    p_train <- stats::predict(learner$forest, data.frame(X_train))
    p_test <- stats::predict(learner$forest, data.frame(X_test))
    eval_points <- sort(unique(y_train[, 1]))
    eval_points <- eval_points[eval_points >= min(y_test[, 1])]
    eval_points <- eval_points[eval_points < max(y_test[, 1])]
    pos <- prodlim::sindex(
      jump.times = p_test$unique.death.times,
      eval.times = eval_points
    )

    integrated_brier_score <- ibrier(
      Surv(y_test),
      t(p_test$survival[, pos]),
      eval_points
    )[1, 1]

    train_predictions <- rowSums(p_train$chf)
    test_predictions <- rowSums(p_test$chf)
    concordance <- unname(rcorr.cens(
      -test_predictions,
      Surv(y_test)
    )[1])

    list(
      concordance = concordance,
      ibs = integrated_brier_score,
      params = learner$paramvalues,
      selected_variables = NA,
      train_predictions = train_predictions,
      test_predictions = test_predictions
    )
  } else if (model %in% c("ridge", "glmnet_favor")) {
    set.seed(seed)
    alpha <- 1
    if (model == "ridge") {
      alpha <- 0
    }
    colnames(y_train) <- c("time", "status")
    colnames(y_test) <- c("time", "status")
    nFolds <- 5
    foldid <- sample(rep(seq(nFolds), length.out = nrow(X_train)))
    penalty_factors <- rep(1, ncol(X_train))
    if (grepl("favor", model) & sum(grepl("clin", colnames(X_train))) < ncol(X_train)) {
      # If model is fit with favoring and we have enough features
      # that this is feasible computationally,
      # fit clinical features unpenalized.
      penalty_factors[grep("clin", colnames(X_train))] <- 0
    }
    if (ncol(X_train) > 1) {
      tic()
      learner <- cv.glmnet(
        x = X_train,
        y = y_train,
        type.measure = "C",
        family = "cox",
        nfolds = nFolds,
        foldid = foldid,
        alpha = alpha,
        penalty.factor = penalty_factors
      )
      exectime <- toc()
      exectime <- exectime$toc - exectime$tic
      selected_variables <- rownames(extract.coef(learner, lambda = "lambda.min"))
      params <- c(lambda.min = learner$lambda.min)
    } else if (
      ncol(X_train) == 1
    ) {
      tic()
      learner <- bigGlm(
        x = X_train,
        y = y_train,
        family = "cox"
      )
      exectime <- toc()
      exectime <- exectime$toc - exectime$tic
      selected_variables <- NA
      params <- NA
    }

    train_predictions <- predict(learner,
      newx = X_train,
      # Use minimum lambda for all `glmnet` related models.
      s = "lambda.min", arg = "response"
    )[, 1]
    test_predictions <- predict(learner,
      newx = X_test,
      # Use minimum lambda for all `glmnet` related models.
      s = "lambda.min", arg = "response"
    )[, 1]

    concordance <- unname(rcorr.cens(
      -test_predictions,
      Surv(y_test)
    )[1])
    return(list(
      concordance = concordance,
      ibs = NA,
      params = params,
      selected_variables = selected_variables,
      train_predictions = train_predictions,
      test_predictions = test_predictions
    ))
  } else if (model == "prioritylasso") {
    set.seed(seed)
    nFolds <- 5
    foldid <- sample(rep(seq(nFolds), length.out = nrow(X_train)))
    colnames(y_train) <- c("time", "stop")
    colnames(y_test) <- c("time", "stop")
    tic()
    learner <- prioritylasso(
      X_train,
      Surv(y_train[, 1], y_train[, 2]),
      blocks = blocks,
      nfolds = 5,
      cvoffset = TRUE,
      cvoffsetnfolds = 5,
      family = "cox",
      foldid = foldid,
      type.measure = "deviance",
      block1.penalization = TRUE,
      lambda.type = "lambda.min"
    )
    exectime <- toc()
    exectime <- exectime$toc - exectime$tic
    selected_variables <- names(which(learner$coefficients != 0))
    params <- c(lambda.min = learner$lambda.min)
    # In case there are very strong correlations within clinical data,
    # this can lead to some coefficients being `NA` as prioritylasso
    # favoring fits the first (i.e., clinical) block unpenalized.
    # Thus, we subset to the non-zero (and thus, non-NA) coefficients
    # to calculate the predictions.
    if (length(selected_variables) > 1) {
      train_predictions <- exp(X_train[, which(learner$coefficients != 0)] %*% learner$coefficients[which(learner$coefficients != 0)])[, 1]
      test_predictions <- exp(X_test[, which(learner$coefficients != 0)] %*% learner$coefficients[which(learner$coefficients != 0)])[, 1]
    } else {
      train_predictions <- exp(
        X_train[, which(learner$coefficients != 0)] * learner$coefficients[which(learner$coefficients != 0)]
      )
      test_predictions <- exp(
        X_test[, which(learner$coefficients != 0)] * learner$coefficients[which(learner$coefficients != 0)]
      )
    }

    concordance <- unname(rcorr.cens(
      -test_predictions,
      Surv(y_test)
    )[1])
    return(
      list(
        concordance = concordance,
        ibs = NA,
        params = params,
        selected_variables = selected_variables,
        train_predictions = train_predictions,
        test_predictions = test_predictions
      )
    )
  } else if (model == "ranger") {
    set.seed(seed)
    colnames(y_train) <- c("time", "status")
    colnames(y_test) <- c("time", "status")
    tic()
    learner <- tuneRanger::tuneMtryFast(
      dependent.variable.name = "time",
      status.variable.name = "status",
      data = cbind(y_train, X_train),
      doBest = TRUE,
      trace = FALSE,
      plot = FALSE,
      seed = seed
    )
    exectime <- toc()
    exectime <- exectime$toc - exectime$tic
    selected_variables <- NA
    params <- c(mtry = learner$mtry)

    p_train <- predict(learner, data = X_train)
    p_test <- predict(learner, data = X_test)

    eval_points <- sort(unique(y_train[, 1]))
    eval_points <- eval_points[eval_points >= min(y_test[, 1])]
    eval_points <- eval_points[eval_points < max(y_test[, 1])]
    pos <- prodlim::sindex(
      jump.times = p_test$unique.death.times,
      eval.times = eval_points
    )

    integrated_brier_score <- ibrier(
      Surv(y_test),
      t(p_test$survival[, pos]),
      eval_points
    )[1, 1]

    train_predictions <- rowSums(p_train$chf)
    test_predictions <- rowSums(p_test$chf)
    concordance <- unname(rcorr.cens(
      -test_predictions,
      Surv(y_test)
    )[1])

    list(
      concordance = concordance,
      ibs = integrated_brier_score,
      params = params,
      selected_variables = NA,
      train_predictions = train_predictions,
      test_predictions = test_predictions
    )
  } else if (model == "clinical_ridge") {
    set.seed(seed)
    clinical_col_ix <- grep("clin", colnames(X_train))
    colnames(y_train) <- c("time", "status")
    colnames(y_test) <- c("time", "status")
    nFolds <- 5
    foldid <- sample(rep(seq(nFolds), length.out = nrow(X_train)))
    tic()
    learner <- cv.glmnet(
      x = X_train[, clinical_col_ix],
      y = y_train,
      type.measure = "C",
      family = "cox",
      nfolds = nFolds,
      foldid = foldid,
      alpha = 0
    )
    exectime <- toc()
    exectime <- exectime$toc - exectime$tic
    selected_variables <- rownames(extract.coef(learner))
    params <- c(lambda.min = learner$lambda.min)


    train_predictions <- predict(learner,
      newx = X_train[, clinical_col_ix],
      # Use minimum lambda for all `glmnet` related models.
      s = "lambda.min", arg = "response"
    )[, 1]
    test_predictions <- predict(learner,
      newx = X_test[, clinical_col_ix],
      # Use minimum lambda for all `glmnet` related models.
      s = "lambda.min", arg = "response"
    )[, 1]

    concordance <- unname(rcorr.cens(
      -test_predictions,
      Surv(y_test)
    )[1])
    eval_points <- sort(unique(y_train[, 1]))
    eval_points <- eval_points[eval_points >= min(y_test[, 1])]
    eval_points <- eval_points[eval_points < max(y_test[, 1])]

    cox_helper <- coxph(
      Surv(time, status) ~ .,
      data = cbind(y_train, data.frame(X_train[, colnames(X_train) %in% selected_variables])),
      init = extract.coef(learner, lambda = "lambda.min")[, 1],
      iter.max = 0,
      x = TRUE,
      ties = "breslow"
    )

    survival_probabilities <- pec::predictSurvProb(cox_helper,
      newdata = data.frame(X_test[, colnames(X_train) %in% selected_variables]),
      times = eval_points
    )

    integrated_brier_score <- ibrier(
      Surv(y_test),
      t(survival_probabilities),
      eval_points
    )[1, 1]

    list(
      concordance = concordance,
      ibs = integrated_brier_score,
      params = params,
      selected_variables = selected_variables,
      train_predictions = train_predictions,
      test_predictions = test_predictions
    )
  }
}
```
