## Supplementary material for "Clinically driven knowledge distillation for sparsifying high-dimensional multi-omics survival models": Code: reproduce_figure_1.R

```
library(here)
library(rjson)
library(readr)
library(dplyr)
library(ggplot2)
library(tidyr)
library(ggpubfigs)

config <- rjson::fromJSON(
  file = here::here("config", "config.json")
)
plt_frame <- data.frame()

for (cancer in config$cancers) {
  plt_frame <- rbind(
    plt_frame,
    data.frame(
      model = "Clinical Cox PH",
      cancer = cancer,
      concordance = sapply(
        readRDS(here::here("data", "benchmarks", cancer, "clinical_ridge_10_stratified_all.rds")),
        function(x) x$concordance
      ),
      type = "Clinical-only"
    )
  )

  plt_frame <- rbind(
    plt_frame,
    data.frame(
      model = "BF",
      cancer = cancer,
      concordance = sapply(
        readRDS(here::here("data", "benchmarks", cancer, "blockforest_10_stratified_all.rds")),
        function(x) x$concordance
      ),
      type = "Teacher"
    )
  )

  plt_frame <- rbind(
    plt_frame,
    data.frame(
      model = "RBF",
      cancer = cancer,
      concordance = sapply(
        readRDS(here::here("data", "benchmarks", cancer, "randomblock_favoring_10_stratified_all.rds")),
        function(x) x$concordance
      ),
      type = "Teacher"
    )
  )

  plt_frame <- rbind(
    plt_frame,
    data.frame(
      model = "KD favoring (RBF)",
      cancer = cancer,
      concordance = unlist(sapply(
        readRDS(here::here("data", "benchmarks", cancer, "randomblock_favoring_distilled_fidelity_with_intercept_all.rds")),
        function(x) x[[4]]
      ) %>% replace_na(
        0.5
      )),
      type = "Distillation"
    )
  )

  plt_frame <- rbind(
    plt_frame,
    data.frame(
      model = "KD favoring (HSAE)",
      cancer = cancer,
      concordance = unlist(sapply(
        readRDS(here::here("data", "benchmarks", cancer, "hierarchical_sae_distilled_fidelity_with_intercept_all.rds")),
        function(x) x[[4]]
      ) %>% replace_na(
        0.5
      )),
      type = "Distillation"
    )
  )

  plt_frame <- rbind(
    plt_frame,
    data.frame(
      model = "KD favoring (BF)",
      cancer = cancer,
      concordance = unlist(sapply(
        readRDS(here::here("data", "benchmarks", cancer, "blockforest_distilled_fidelity_with_intercept_all.rds")),
        function(x) x[[4]]
      ) %>% replace_na(
        0.5
      )),
      type = "Distillation"
    )
  )

  plt_frame <- rbind(
    plt_frame,
    (read_csv(here::here("data", "benchmarks", cancer, "hierarchical_sae_scores_all_es.csv"))) %>%
      dplyr::select(model, concordance) %>%
      mutate(cancer = cancer) %>%
      mutate(model = "HSAE") %>%
      mutate(type = "Teacher")
  )
}

plt_frame$model <- factor(plt_frame$model, c(
  "HSAE", "RBF", "BF", "KD favoring (HSAE)", "KD favoring (RBF)", "KD favoring (BF)", "Clinical Cox PH"
))

plt_frame$type <- factor(plt_frame$type, c("Teacher", "Distillation", "Clinical-only"))

plt_frame %>%
  ggplot(aes(x = model, y = concordance, fill = type)) +
  geom_boxplot(outlier.size = 0.5, fatten = 1.5) +
  facet_wrap(~cancer, scales = "free_y") +
  labs(x = "Model", y = "Test concordance") +
  theme_bw(base_size = 8) +
  scale_fill_manual(values = friendly_pal("ito_seven")[c(2, 1, 3)], name = "Model type") +
  theme(axis.text.x = element_text(angle = 90)) -> p

p

ggsave(filename = here::here("figures", "fig-1.pdf"), device = "pdf", dpi = 600, height = 9, width = 17.1425, units = "cm")
```
