## Supplementary material for "Clinically driven knowledge distillation for sparsifying high-dimensional multi-omics survival models": Code: reproduce_figure_2.R

```
library(here)
library(rjson)
library(readr)
library(dplyr)
library(ggplot2)
library(tidyr)
library(ggpubfigs)

config <- rjson::fromJSON(
  file = here::here("config", "config.json")
)
plt_frame <- data.frame()

for (cancer in config$cancers) {
  plt_frame <- rbind(
    plt_frame,
    data.frame(
      model = "Lasso favoring",
      cancer = cancer,
      concordance = sapply(
        readRDS(here::here("data", "benchmarks", cancer, "glmnet_favor_10_stratified_all.rds")),
        function(x) x$concordance
      ),
      type = "Regression"
    )
  )

  plt_frame <- rbind(
    plt_frame,
    data.frame(
      model = "prioritylasso",
      cancer = cancer,
      concordance = sapply(
        readRDS(here::here("data", "benchmarks", cancer, "prioritylasso_10_stratified_all.rds")),
        function(x) x$concordance
      ),
      type = "Regression"
    )
  )

  plt_frame <- rbind(
    plt_frame,
    data.frame(
      model = "KD favoring (RBF)",
      cancer = cancer,
      concordance = unlist(sapply(
        readRDS(here::here("data", "benchmarks", cancer, "randomblock_favoring_distilled_fidelity_with_intercept_all.rds")),
        function(x) x[[4]]
      ) %>% replace_na(
        0.5
      )),
      type = "Distillation"
    )
  )

  plt_frame <- rbind(
    plt_frame,
    data.frame(
      model = "KD favoring (HSAE)",
      cancer = cancer,
      concordance = unlist(sapply(
        readRDS(here::here("data", "benchmarks", cancer, "hierarchical_sae_distilled_fidelity_with_intercept_all.rds")),
        function(x) x[[4]]
      ) %>% replace_na(
        0.5
      )),
      type = "Distillation"
    )
  )

  plt_frame <- rbind(
    plt_frame,
    data.frame(
      model = "Clinical Cox PH",
      cancer = cancer,
      concordance = unlist(sapply(
        readRDS(here::here("data", "benchmarks", cancer, "clinical_ridge_10_stratified_all.rds")),
        function(x) x$concordance
      ) %>% replace_na(
        0.5
      )),
      type = "Clinical-only"
    )
  )

  plt_frame <- rbind(
    plt_frame,
    data.frame(
      model = "KD favoring (BF)",
      cancer = cancer,
      concordance = unlist(sapply(
        readRDS(here::here("data", "benchmarks", cancer, "blockforest_distilled_fidelity_with_intercept_all.rds")),
        function(x) x[[4]]
      ) %>% replace_na(
        0.5
      )),
      type = "Distillation"
    )
  )
}


plt_frame$model <- factor(plt_frame$model, levels = c(
  "KD favoring (HSAE)", "KD favoring (RBF)", "KD favoring (BF)",
  "prioritylasso", "Lasso favoring",
  "Clinical Cox PH"
))
plt_frame$type <- factor(plt_frame$type, levels = c(
  "Distillation", "Regression", "Clinical-only"
))

plt_frame %>%
  group_by(model, cancer) %>%
  summarise(mean_concordance = mean(concordance), .groups = "keep") -> tmp


p_vals <- data.frame(
  Model = unique(plt_frame$model)[-grep("Clinical Cox PH", unique(plt_frame$model), fixed = TRUE)],
  p.val = sapply(
    unique(plt_frame$model)[-grep("Clinical Cox PH", unique(plt_frame$model), fixed = TRUE)],
    function(x) {
      wilcox.test(
        x = tmp %>% filter(model == "Clinical Cox PH") %>% pull(mean_concordance),
        y = tmp %>% filter(model == x) %>% pull(mean_concordance),
        paired = TRUE,
        mu = 0,
        alternative = "less",
        digits.rank = 7
      )$p.value
    }
  )
)

library(ggsignif)
plt_frame %>%
  ggplot(aes(x = model, y = concordance, fill = type)) +
  geom_boxplot(outlier.size = 0.5, fatten = 1.5) +
  theme_bw(base_size = 8) +
  geom_signif(
    annotations = paste0("p = ", round(p_vals$p.val[which(p_vals$p.val < 0.01)], 4)),
    xmin = p_vals$Model[which(p_vals$p.val < 0.01)],
    xmax = rep("Clinical Cox PH", sum(p_vals$p.val < 0.01)),
    y_position = c(0.9675, 1, 0.935),
    tip_length = 0,
    textsize = 3.88 / 2.5
  ) +
  scale_fill_manual(values = friendly_pal("ito_seven")[c(1, 4, 3)], name = "Model type") +
  theme(axis.text.x = element_text(angle = 90)) +
  labs(x = "", y = "Test concordance") -> p

p
ggsave(filename = here::here("figures", "fig-2.pdf"), device = "pdf", dpi = 600, height = 7, width = 9, units = "cm")
```
