## Supplementary material for "Clinically driven knowledge distillation for sparsifying high-dimensional multi-omics survival models": Code: reproduce_figure_s2.R

```
library(here)
library(rjson)
library(readr)
library(dplyr)
library(ggplot2)
library(tidyr)

config <- rjson::fromJSON(
  file = here::here("config", "config.json")
)
plt_frame <- data.frame()

for (cancer in config$cancers) {
  plt_frame <- rbind(
    plt_frame,
    data.frame(
      model = "KD favoring (RBF)",
      cancer = cancer,
      train_correlation = sapply(readRDS(here::here("data", "benchmarks", cancer, "randomblock_favoring_distilled_fidelity_with_intercept_all.rds")), function(x) x[[6]][[1]]),
      test_correlation = sapply(readRDS(here::here("data", "benchmarks", cancer, "randomblock_favoring_distilled_fidelity_with_intercept_all.rds")), function(x) x[[6]][[2]])
    )
  )

  plt_frame <- rbind(
    plt_frame,
    data.frame(
      model = "KD favoring (HSAE)",
      cancer = cancer,
      train_correlation = sapply(readRDS(here::here("data", "benchmarks", cancer, "hierarchical_sae_distilled_fidelity_with_intercept_all.rds")), function(x) x[[6]][[1]]),
      test_correlation = sapply(readRDS(here::here("data", "benchmarks", cancer, "hierarchical_sae_distilled_fidelity_with_intercept_all.rds")), function(x) x[[6]][[2]])
    )
  )

  plt_frame <- rbind(
    plt_frame,
    data.frame(
      model = "KD favoring (BF)",
      cancer = cancer,
      train_correlation = sapply(readRDS(here::here("data", "benchmarks", cancer, "blockforest_distilled_fidelity_with_intercept_all.rds")), function(x) x[[6]][[1]]),
      test_correlation = sapply(readRDS(here::here("data", "benchmarks", cancer, "blockforest_distilled_fidelity_with_intercept_all.rds")), function(x) x[[6]][[2]])
    )
  )
}

plt_frame$model <- factor(plt_frame$model, c("KD favoring (HSAE)", "KD favoring (RBF)", "KD favoring (BF)"))

tmpo %>% ggplot(aes(x = model, y = test_correlation)) +
  geom_boxplot(outlier.size = 0.5, fatten = 1.5) +
  facet_wrap(~cancer, scales = "free_y") +
  labs(x = "Model", y = "Test rank correlation") +
  theme_bw(base_size = 8) +
  theme(legend.title = element_blank()) +
  theme(axis.text.x = element_text(angle = 90)) -> p

p
ggsave(filename = here::here("figures", "fig-s2.pdf"), device = "pdf", dpi = 600, height = 12, width = 17.1425, units = "cm")
```
