## Supplementary material for "Clinically driven knowledge distillation for sparsifying high-dimensional multi-omics survival models": Code: reproduce_table_1.R

```
library(here)
library(rjson)
library(readr)
library(dplyr)
library(ggplot2)
library(tidyr)

config <- rjson::fromJSON(
  file = here::here("config", "config.json")
)
plt_frame <- data.frame()

for (cancer in config$cancers) {
  plt_frame <- rbind(
    plt_frame,
    data.frame(
      model = "BlockForest",
      cancer = cancer,
      concordance = sapply(
        readRDS(here::here("data", "benchmarks", cancer, "blockforest_10_stratified_all.rds")),
        function(x) x$concordance
      ),
      train_correlation = NA,
      test_correlation = NA,
      total = "79329.8 (3815.2)"
    )
  )

  plt_frame <- rbind(
    plt_frame,
    data.frame(
      model = "RandomBlock favoring",
      cancer = cancer,
      concordance = sapply(
        readRDS(here::here("data", "benchmarks", cancer, "randomblock_favoring_10_stratified_all.rds")),
        function(x) x$concordance
      ),
      train_correlation = NA,
      test_correlation = NA,
      total = "79329.8 (3815.2)"
    )
  )

  plt_frame <- rbind(
    plt_frame,
    data.frame(
      model = "KD favoring (RBF)",
      cancer = cancer,
      concordance = unlist(sapply(
        readRDS(here::here("data", "benchmarks", cancer, "randomblock_favoring_distilled_fidelity_with_intercept_all.rds")),
        function(x) x[[4]]
      ) %>% replace_na(
        0.5
      )),
      total = sapply(readRDS(here::here("data", "benchmarks", cancer, "randomblock_favoring_distilled_fidelity_with_intercept_all.rds")), function(x) length(x[[5]])),
      train_correlation = sapply(readRDS(here::here("data", "benchmarks", cancer, "randomblock_favoring_distilled_fidelity_with_intercept_all.rds")), function(x) x[[6]][[1]]),
      test_correlation = sapply(readRDS(here::here("data", "benchmarks", cancer, "randomblock_favoring_distilled_fidelity_with_intercept_all.rds")), function(x) x[[6]][[2]])
    )
  )

  plt_frame <- rbind(
    plt_frame,
    data.frame(
      model = "KD (RBF)",
      cancer = cancer,
      concordance = unlist(sapply(
        readRDS(here::here("data", "benchmarks", cancer, "randomblock_favoring_distilled_fidelity_with_intercept_all.rds")),
        function(x) x[[1]]
      ) %>% replace_na(
        0.5
      )),
      total = sapply(readRDS(here::here("data", "benchmarks", cancer, "randomblock_favoring_distilled_fidelity_with_intercept_all.rds")), function(x) length(x[[2]])),
      train_correlation = sapply(readRDS(here::here("data", "benchmarks", cancer, "randomblock_favoring_distilled_fidelity_with_intercept_all.rds")), function(x) x[[3]][[1]]),
      test_correlation = sapply(readRDS(here::here("data", "benchmarks", cancer, "randomblock_favoring_distilled_fidelity_with_intercept_all.rds")), function(x) x[[3]][[2]])
    )
  )

  plt_frame <- rbind(
    plt_frame,
    data.frame(
      model = "KD favoring (HSAE)",
      cancer = cancer,
      concordance = unlist(sapply(
        readRDS(here::here("data", "benchmarks", cancer, "hierarchical_sae_distilled_fidelity_with_intercept_all.rds")),
        function(x) x[[4]]
      ) %>% replace_na(
        0.5
      )),
      total = sapply(readRDS(here::here("data", "benchmarks", cancer, "hierarchical_sae_distilled_fidelity_with_intercept_all.rds")), function(x) length(x[[5]])),
      train_correlation = sapply(readRDS(here::here("data", "benchmarks", cancer, "hierarchical_sae_distilled_fidelity_with_intercept_all.rds")), function(x) x[[6]][[1]]),
      test_correlation = sapply(readRDS(here::here("data", "benchmarks", cancer, "hierarchical_sae_distilled_fidelity_with_intercept_all.rds")), function(x) x[[6]][[2]])
    )
  )

  plt_frame <- rbind(
    plt_frame,
    data.frame(
      model = "KD (HSAE)",
      cancer = cancer,
      concordance = unlist(sapply(
        readRDS(here::here("data", "benchmarks", cancer, "hierarchical_sae_distilled_fidelity_with_intercept_all.rds")),
        function(x) x[[1]]
      ) %>% replace_na(
        0.5
      )),
      total = sapply(readRDS(here::here("data", "benchmarks", cancer, "hierarchical_sae_distilled_fidelity_with_intercept_all.rds")), function(x) length(x[[2]])),
      train_correlation = sapply(readRDS(here::here("data", "benchmarks", cancer, "hierarchical_sae_distilled_fidelity_with_intercept_all.rds")), function(x) x[[3]][[1]]),
      test_correlation = sapply(readRDS(here::here("data", "benchmarks", cancer, "hierarchical_sae_distilled_fidelity_with_intercept_all.rds")), function(x) x[[3]][[2]])
    )
  )

  plt_frame <- rbind(
    plt_frame,
    data.frame(
      model = "KD favoring (BF)",
      cancer = cancer,
      concordance = unlist(sapply(
        readRDS(here::here("data", "benchmarks", cancer, "blockforest_distilled_fidelity_with_intercept_all.rds")),
        function(x) x[[4]]
      ) %>% replace_na(
        0.5
      )),
      total = sapply(readRDS(here::here("data", "benchmarks", cancer, "blockforest_distilled_fidelity_with_intercept_all.rds")), function(x) length(x[[5]])),
      train_correlation = sapply(readRDS(here::here("data", "benchmarks", cancer, "blockforest_distilled_fidelity_with_intercept_all.rds")), function(x) x[[6]][[1]]),
      test_correlation = sapply(readRDS(here::here("data", "benchmarks", cancer, "blockforest_distilled_fidelity_with_intercept_all.rds")), function(x) x[[6]][[2]])
    )
  )

  plt_frame <- rbind(
    plt_frame,
    data.frame(
      model = "KD (BF)",
      cancer = cancer,
      concordance = unlist(sapply(
        readRDS(here::here("data", "benchmarks", cancer, "blockforest_distilled_fidelity_with_intercept_all.rds")),
        function(x) x[[1]]
      ) %>% replace_na(
        0.5
      )),
      total = sapply(readRDS(here::here("data", "benchmarks", cancer, "blockforest_distilled_fidelity_with_intercept_all.rds")), function(x) length(x[[2]])),
      train_correlation = sapply(readRDS(here::here("data", "benchmarks", cancer, "blockforest_distilled_fidelity_with_intercept_all.rds")), function(x) x[[3]][[1]]),
      test_correlation = sapply(readRDS(here::here("data", "benchmarks", cancer, "blockforest_distilled_fidelity_with_intercept_all.rds")), function(x) x[[3]][[2]])
    )
  )
  plt_frame <- rbind(
    plt_frame,
    (read_csv(here::here("data", "benchmarks", cancer, "hierarchical_sae_scores_all_es.csv"))) %>%
      dplyr::select(model, concordance) %>%
      mutate(total = "79329.8 (3815.2)") %>%
      mutate(cancer = cancer) %>%
      mutate(model = "HSAE") %>%
      mutate(type = "Teacher")
  )
}

plt_frame %>%
  group_by(model) %>%
  summarise(
    mean_concordance = mean(concordance), sd_concordance = sd(concordance, na.rm = TRUE),
    mean_train_r2 = mean(train_correlation, na.rm = TRUE), sd_train_r2 = sd(train_correlation, na.rm = TRUE),
    mean_test_r2 = mean(test_correlation, na.rm = TRUE), sd_test_r2 = sd(test_correlation, na.rm = TRUE),
    mean_coefs = mean(total, na.rm = TRUE), sd_coefs = sd(total, na.rm = TRUE)
  ) %>%
  replace(is.na(.), 0) %>%
  mutate(across(2:9, round, 3)) %>%
  mutate(Concordance = paste0(mean_concordance, " (", sd_concordance, ")")) %>%
  mutate(`Train R2` = paste0(mean_train_r2, " (", sd_train_r2, ")")) %>%
  mutate(`Test R2` = paste0(mean_test_r2, " (", sd_test_r2, ")")) %>%
  mutate(`Molecular features used` = paste0(mean_coefs, " (", sd_coefs, ")")) %>%
  select(model, Concordance, `Train R2`, `Test R2`, `Molecular features used`) -> tmp

tmp$Student <- c("None", "None", "Lasso", "Lasso", "Lasso", "Lasso favoring clinical variables", "Lasso favoring clinical variables", "Lasso favoring clinical variables", "None")
tmp <- tmp[, c(1, ncol(tmp), 2:5)]
colnames(tmp)[1:2] <- c("Teacher", "Student")
tmp$Teacher <- c(
  "BF",
  "HSAE",
  "BF",
  "HSAE",
  "RBF",
  "BF",
  "HSAE",
  "RBF",
  "RBF"
)

tmp %>% write_csv(
  here::here(
    "tables", "table_1.csv"
  )
)
```
