## Supplementary material for "Clinically driven knowledge distillation for sparsifying high-dimensional multi-omics survival models": Code: reproduce_table_2.R

```
library(here)
library(rjson)
library(readr)
library(dplyr)
library(ggplot2)
library(tidyr)

config <- rjson::fromJSON(
  file = here::here("config", "config.json")
)
plt_frame <- data.frame()

for (cancer in config$cancers) {
  plt_frame <- rbind(
    plt_frame,
    data.frame(
      model = "KD favoring (RBF)",
      cancer = cancer,
      split = 1:10,
      Clinical = sapply(readRDS(here::here("data", "benchmarks", cancer, "randomblock_favoring_distilled_fidelity_with_intercept_all.rds")), function(x) sum(grepl("clinical", x[[5]]))),
      GEX = sapply(readRDS(here::here("data", "benchmarks", cancer, "randomblock_favoring_distilled_fidelity_with_intercept_all.rds")), function(x) sum(grepl("gex", x[[5]]))),
      Mutation = sapply(readRDS(here::here("data", "benchmarks", cancer, "randomblock_favoring_distilled_fidelity_with_intercept_all.rds")), function(x) sum(grepl("mut", x[[5]]))),
      Methylation = sapply(readRDS(here::here("data", "benchmarks", cancer, "randomblock_favoring_distilled_fidelity_with_intercept_all.rds")), function(x) sum(grepl("meth", x[[5]]))),
      RPPA = sapply(readRDS(here::here("data", "benchmarks", cancer, "randomblock_favoring_distilled_fidelity_with_intercept_all.rds")), function(x) sum(grepl("rppa", x[[5]]))),
      miRNA = sapply(readRDS(here::here("data", "benchmarks", cancer, "randomblock_favoring_distilled_fidelity_with_intercept_all.rds")), function(x) sum(grepl("mirna", x[[5]]))),
      CNV = sapply(readRDS(here::here("data", "benchmarks", cancer, "randomblock_favoring_distilled_fidelity_with_intercept_all.rds")), function(x) sum(grepl("cnv", x[[5]])))
    )
  )

  plt_frame <- rbind(
    plt_frame,
    data.frame(
      model = "KD favoring (BF)",
      cancer = cancer,
      split = 1:10,
      Clinical = sapply(readRDS(here::here("data", "benchmarks", cancer, "blockforest_distilled_fidelity_with_intercept_all.rds")), function(x) sum(grepl("clinical", x[[5]]))),
      GEX = sapply(readRDS(here::here("data", "benchmarks", cancer, "blockforest_distilled_fidelity_with_intercept_all.rds")), function(x) sum(grepl("gex", x[[5]]))),
      Mutation = sapply(readRDS(here::here("data", "benchmarks", cancer, "blockforest_distilled_fidelity_with_intercept_all.rds")), function(x) sum(grepl("mut", x[[5]]))),
      Methylation = sapply(readRDS(here::here("data", "benchmarks", cancer, "blockforest_distilled_fidelity_with_intercept_all.rds")), function(x) sum(grepl("meth", x[[5]]))),
      RPPA = sapply(readRDS(here::here("data", "benchmarks", cancer, "blockforest_distilled_fidelity_with_intercept_all.rds")), function(x) sum(grepl("rppa", x[[5]]))),
      miRNA = sapply(readRDS(here::here("data", "benchmarks", cancer, "blockforest_distilled_fidelity_with_intercept_all.rds")), function(x) sum(grepl("mirna", x[[5]]))),
      CNV = sapply(readRDS(here::here("data", "benchmarks", cancer, "blockforest_distilled_fidelity_with_intercept_all.rds")), function(x) sum(grepl("cnv", x[[5]])))
    )
  )

  plt_frame <- rbind(
    plt_frame,
    data.frame(
      model = "KD favoring (HSAE)",
      cancer = cancer,
      split = 1:10,
      Clinical = sapply(readRDS(here::here("data", "benchmarks", cancer, "hierarchical_sae_distilled_fidelity_with_intercept_all.rds")), function(x) sum(grepl("clinical", x[[5]]))),
      GEX = sapply(readRDS(here::here("data", "benchmarks", cancer, "hierarchical_sae_distilled_fidelity_with_intercept_all.rds")), function(x) sum(grepl("gex", x[[5]]))),
      Mutation = sapply(readRDS(here::here("data", "benchmarks", cancer, "hierarchical_sae_distilled_fidelity_with_intercept_all.rds")), function(x) sum(grepl("mut", x[[5]]))),
      Methylation = sapply(readRDS(here::here("data", "benchmarks", cancer, "hierarchical_sae_distilled_fidelity_with_intercept_all.rds")), function(x) sum(grepl("meth", x[[5]]))),
      RPPA = sapply(readRDS(here::here("data", "benchmarks", cancer, "hierarchical_sae_distilled_fidelity_with_intercept_all.rds")), function(x) sum(grepl("rppa", x[[5]]))),
      miRNA = sapply(readRDS(here::here("data", "benchmarks", cancer, "hierarchical_sae_distilled_fidelity_with_intercept_all.rds")), function(x) sum(grepl("mirna", x[[5]]))),
      CNV = sapply(readRDS(here::here("data", "benchmarks", cancer, "hierarchical_sae_distilled_fidelity_with_intercept_all.rds")), function(x) sum(grepl("cnv", x[[5]])))
    )
  )

  plt_frame <- rbind(
    plt_frame,
    data.frame(
      model = "Lasso favoring",
      cancer = cancer,
      split = 1:10,
      Clinical = sapply(readRDS(here::here("data", "benchmarks", cancer, "glmnet_favor_10_stratified_all.rds")), function(x) sum(grepl("clinical", x$selected_variables))),
      GEX = sapply(readRDS(here::here("data", "benchmarks", cancer, "glmnet_favor_10_stratified_all.rds")), function(x) sum(grepl("gex", x$selected_variables))),
      Mutation = sapply(readRDS(here::here("data", "benchmarks", cancer, "glmnet_favor_10_stratified_all.rds")), function(x) sum(grepl("mut", x$selected_variables))),
      Methylation = sapply(readRDS(here::here("data", "benchmarks", cancer, "glmnet_favor_10_stratified_all.rds")), function(x) sum(grepl("meth", x$selected_variables))),
      RPPA = sapply(readRDS(here::here("data", "benchmarks", cancer, "glmnet_favor_10_stratified_all.rds")), function(x) sum(grepl("rppa", x$selected_variables))),
      miRNA = sapply(readRDS(here::here("data", "benchmarks", cancer, "glmnet_favor_10_stratified_all.rds")), function(x) sum(grepl("mirna", x$selected_variables))),
      CNV = sapply(readRDS(here::here("data", "benchmarks", cancer, "glmnet_favor_10_stratified_all.rds")), function(x) sum(grepl("cnv", x$selected_variables)))
    )
  )

  plt_frame <- rbind(
    plt_frame,
    data.frame(
      model = "prioritylasso",
      cancer = cancer,
      split = 1:10,
      Clinical = sapply(readRDS(here::here("data", "benchmarks", cancer, "prioritylasso_10_stratified_all.rds")), function(x) sum(grepl("clinical", x$selected_variables))),
      GEX = sapply(readRDS(here::here("data", "benchmarks", cancer, "prioritylasso_10_stratified_all.rds")), function(x) sum(grepl("gex", x$selected_variables))),
      Mutation = sapply(readRDS(here::here("data", "benchmarks", cancer, "prioritylasso_10_stratified_all.rds")), function(x) sum(grepl("mut", x$selected_variables))),
      Methylation = sapply(readRDS(here::here("data", "benchmarks", cancer, "prioritylasso_10_stratified_all.rds")), function(x) sum(grepl("meth", x$selected_variables))),
      RPPA = sapply(readRDS(here::here("data", "benchmarks", cancer, "prioritylasso_10_stratified_all.rds")), function(x) sum(grepl("rppa", x$selected_variables))),
      miRNA = sapply(readRDS(here::here("data", "benchmarks", cancer, "prioritylasso_10_stratified_all.rds")), function(x) sum(grepl("mirna", x$selected_variables))),
      CNV = sapply(readRDS(here::here("data", "benchmarks", cancer, "prioritylasso_10_stratified_all.rds")), function(x) sum(grepl("cnv", x$selected_variables)))
    )
  )
}

plt_frame %>%
  group_by(model) %>%
  summarise(
    Clinical = paste0(round(mean(Clinical), 1), " (", round(sd(Clinical), 1), ")"),
    GEX = paste0(round(mean(GEX), 1), " (", round(sd(GEX), 1), ")"),
    Mutation = paste0(round(mean(Mutation), 1), " (", round(sd(Mutation), 1), ")"),
    Methylation = paste0(round(mean(Methylation), 1), " (", round(sd(Methylation), 1), ")"),
    RPPA = paste0(round(mean(RPPA), 1), " (", round(sd(RPPA), 1), ")"),
    miRNA = paste0(round(mean(miRNA), 1), " (", round(sd(miRNA), 1), ")"),
    CNV = paste0(round(mean(CNV), 1), " (", round(sd(CNV), 1), ")")
  ) %>%
  write_csv(
    here::here(
      "tables", "table_2.csv"
    )
  )
```
