## Supplementary material for "Clinically driven knowledge distillation for sparsifying high-dimensional multi-omics survival models": Code: reproduce_table_3.R

```
library(here)
library(rjson)
library(readr)
library(dplyr)
library(ggplot2)
library(tidyr)

config <- rjson::fromJSON(
  file = here::here("config", "config.json")
)
plt_frame <- data.frame()

for (cancer in config$cancers) {
  plt_frame <- rbind(
    plt_frame,
    data.frame(
      model = "KD favoring (BF)",
      modalities = "Clinical + GEX",
      cancer = cancer,
      concordance = unlist(sapply(
        readRDS(here::here("data", "benchmarks", cancer, "blockforest_distilled_clinical_gex.rds")),
        function(x) x[[4]]
      ) %>% replace_na(
        0.5
      ))
    )
  )

  plt_frame <- rbind(
    plt_frame,
    data.frame(
      model = "KD favoring (BF)",
      modalities = "Clinical + Methylation",
      cancer = cancer,
      concordance = unlist(sapply(
        readRDS(here::here("data", "benchmarks", cancer, "blockforest_distilled_clinical_meth.rds")),
        function(x) x[[4]]
      ) %>% replace_na(
        0.5
      ))
    )
  )

  plt_frame <- rbind(
    plt_frame,
    data.frame(
      model = "KD favoring (BF)",
      modalities = "Clinical + Mutation",
      cancer = cancer,
      concordance = unlist(sapply(
        readRDS(here::here("data", "benchmarks", cancer, "blockforest_distilled_clinical_mut.rds")),
        function(x) x[[4]]
      ) %>% replace_na(
        0.5
      ))
    )
  )

  plt_frame <- rbind(
    plt_frame,
    data.frame(
      model = "KD favoring (BF)",
      modalities = "Clinical + RPPA",
      cancer = cancer,
      concordance = unlist(sapply(
        readRDS(here::here("data", "benchmarks", cancer, "blockforest_distilled_clinical_rppa.rds")),
        function(x) x[[4]]
      ) %>% replace_na(
        0.5
      ))
    )
  )

  plt_frame <- rbind(
    plt_frame,
    data.frame(
      model = "KD favoring (BF)",
      modalities = "Clinical + miRNA",
      cancer = cancer,
      concordance = unlist(sapply(
        readRDS(here::here("data", "benchmarks", cancer, "blockforest_distilled_clinical_mirna.rds")),
        function(x) x[[4]]
      ) %>% replace_na(
        0.5
      ))
    )
  )

  plt_frame <- rbind(
    plt_frame,
    data.frame(
      model = "KD favoring (BF)",
      modalities = "Clinical + CNV",
      cancer = cancer,
      concordance = unlist(sapply(
        readRDS(here::here("data", "benchmarks", cancer, "blockforest_distilled_clinical_cnv.rds")),
        function(x) x[[4]]
      ) %>% replace_na(
        0.5
      ))
    )
  )

  plt_frame <- rbind(
    plt_frame,
    data.frame(
      model = "KD favoring (RBF)",
      modalities = "Clinical + GEX",
      cancer = cancer,
      concordance = unlist(sapply(
        readRDS(here::here("data", "benchmarks", cancer, "randomblock_favoring_distilled_clinical_gex.rds")),
        function(x) x[[4]]
      ) %>% replace_na(
        0.5
      ))
    )
  )

  plt_frame <- rbind(
    plt_frame,
    data.frame(
      model = "KD favoring (RBF)",
      modalities = "Clinical + Methylation",
      cancer = cancer,
      concordance = unlist(sapply(
        readRDS(here::here("data", "benchmarks", cancer, "randomblock_favoring_distilled_clinical_meth.rds")),
        function(x) x[[4]]
      ) %>% replace_na(
        0.5
      ))
    )
  )

  plt_frame <- rbind(
    plt_frame,
    data.frame(
      model = "KD favoring (RBF)",
      modalities = "Clinical + Mutation",
      cancer = cancer,
      concordance = unlist(sapply(
        readRDS(here::here("data", "benchmarks", cancer, "randomblock_favoring_distilled_clinical_mut.rds")),
        function(x) x[[4]]
      ) %>% replace_na(
        0.5
      ))
    )
  )

  plt_frame <- rbind(
    plt_frame,
    data.frame(
      model = "KD favoring (RBF)",
      modalities = "Clinical + RPPA",
      cancer = cancer,
      concordance = unlist(sapply(
        readRDS(here::here("data", "benchmarks", cancer, "randomblock_favoring_distilled_clinical_rppa.rds")),
        function(x) x[[4]]
      ) %>% replace_na(
        0.5
      ))
    )
  )

  plt_frame <- rbind(
    plt_frame,
    data.frame(
      model = "KD favoring (RBF)",
      modalities = "Clinical + miRNA",
      cancer = cancer,
      concordance = unlist(sapply(
        readRDS(here::here("data", "benchmarks", cancer, "randomblock_favoring_distilled_clinical_mirna.rds")),
        function(x) x[[4]]
      ) %>% replace_na(
        0.5
      ))
    )
  )

  plt_frame <- rbind(
    plt_frame,
    data.frame(
      model = "KD favoring (RBF)",
      modalities = "Clinical + CNV",
      cancer = cancer,
      concordance = unlist(sapply(
        readRDS(here::here("data", "benchmarks", cancer, "randomblock_favoring_distilled_clinical_cnv.rds")),
        function(x) x[[4]]
      ) %>% replace_na(
        0.5
      ))
    )
  )


  plt_frame <- rbind(
    plt_frame,
    data.frame(
      model = "KD favoring (HSAE)",
      modalities = "Clinical + GEX",
      cancer = cancer,
      concordance = unlist(sapply(
        readRDS(here::here("data", "benchmarks", cancer, "hierarchical_sae_distilled_clinical_gex.rds")),
        function(x) x[[4]]
      ) %>% replace_na(
        0.5
      ))
    )
  )

  plt_frame <- rbind(
    plt_frame,
    data.frame(
      model = "KD favoring (HSAE)",
      modalities = "Clinical + Methylation",
      cancer = cancer,
      concordance = unlist(sapply(
        readRDS(here::here("data", "benchmarks", cancer, "hierarchical_sae_distilled_clinical_meth.rds")),
        function(x) x[[4]]
      ) %>% replace_na(
        0.5
      ))
    )
  )

  plt_frame <- rbind(
    plt_frame,
    data.frame(
      model = "KD favoring (HSAE)",
      modalities = "Clinical + Mutation",
      cancer = cancer,
      concordance = unlist(sapply(
        readRDS(here::here("data", "benchmarks", cancer, "hierarchical_sae_distilled_clinical_mut.rds")),
        function(x) x[[4]]
      ) %>% replace_na(
        0.5
      ))
    )
  )

  plt_frame <- rbind(
    plt_frame,
    data.frame(
      model = "KD favoring (HSAE)",
      modalities = "Clinical + RPPA",
      cancer = cancer,
      concordance = unlist(sapply(
        readRDS(here::here("data", "benchmarks", cancer, "hierarchical_sae_distilled_clinical_rppa.rds")),
        function(x) x[[4]]
      ) %>% replace_na(
        0.5
      ))
    )
  )

  plt_frame <- rbind(
    plt_frame,
    data.frame(
      model = "KD favoring (HSAE)",
      modalities = "Clinical + miRNA",
      cancer = cancer,
      concordance = unlist(sapply(
        readRDS(here::here("data", "benchmarks", cancer, "hierarchical_sae_distilled_clinical_mirna.rds")),
        function(x) x[[4]]
      ) %>% replace_na(
        0.5
      ))
    )
  )

  plt_frame <- rbind(
    plt_frame,
    data.frame(
      model = "KD favoring (HSAE)",
      modalities = "Clinical + CNV",
      cancer = cancer,
      concordance = unlist(sapply(
        readRDS(here::here("data", "benchmarks", cancer, "hierarchical_sae_distilled_clinical_cnv.rds")),
        function(x) x[[4]]
      ) %>% replace_na(
        0.5
      ))
    )
  )

  plt_frame <- rbind(
    plt_frame,
    data.frame(
      model = "prioritylasso",
      modalities = "Clinical + GEX",
      cancer = cancer,
      concordance = unlist(sapply(
        readRDS(here::here("data", "benchmarks", cancer, "prioritylasso_10_stratified_clinical_gex.rds")),
        function(x) x$concordance
      ) %>% replace_na(
        0.5
      ))
    )
  )

  plt_frame <- rbind(
    plt_frame,
    data.frame(
      model = "prioritylasso",
      modalities = "Clinical + Mutation",
      cancer = cancer,
      concordance = unlist(sapply(
        readRDS(here::here("data", "benchmarks", cancer, "prioritylasso_10_stratified_clinical_mut.rds")),
        function(x) x$concordance
      ) %>% replace_na(
        0.5
      ))
    )
  )

  plt_frame <- rbind(
    plt_frame,
    data.frame(
      model = "prioritylasso",
      modalities = "Clinical + Methylation",
      cancer = cancer,
      concordance = unlist(sapply(
        readRDS(here::here("data", "benchmarks", cancer, "prioritylasso_10_stratified_clinical_meth.rds")),
        function(x) x$concordance
      ) %>% replace_na(
        0.5
      ))
    )
  )

  plt_frame <- rbind(
    plt_frame,
    data.frame(
      model = "prioritylasso",
      modalities = "Clinical + RPPA",
      cancer = cancer,
      concordance = unlist(sapply(
        readRDS(here::here("data", "benchmarks", cancer, "prioritylasso_10_stratified_clinical_rppa.rds")),
        function(x) x$concordance
      ) %>% replace_na(
        0.5
      ))
    )
  )

  plt_frame <- rbind(
    plt_frame,
    data.frame(
      model = "prioritylasso",
      modalities = "Clinical + miRNA",
      cancer = cancer,
      concordance = unlist(sapply(
        readRDS(here::here("data", "benchmarks", cancer, "prioritylasso_10_stratified_clinical_mirna.rds")),
        function(x) x$concordance
      ) %>% replace_na(
        0.5
      ))
    )
  )

  plt_frame <- rbind(
    plt_frame,
    data.frame(
      model = "prioritylasso",
      modalities = "Clinical + CNV",
      cancer = cancer,
      concordance = unlist(sapply(
        readRDS(here::here("data", "benchmarks", cancer, "prioritylasso_10_stratified_clinical_cnv.rds")),
        function(x) x$concordance
      ) %>% replace_na(
        0.5
      ))
    )
  )

  plt_frame <- rbind(
    plt_frame,
    data.frame(
      model = "Lasso favoring",
      modalities = "Clinical + GEX",
      cancer = cancer,
      concordance = unlist(sapply(
        readRDS(here::here("data", "benchmarks", cancer, "glmnet_favor_10_stratified_clinical_gex.rds")),
        function(x) x$concordance
      ) %>% replace_na(
        0.5
      ))
    )
  )

  plt_frame <- rbind(
    plt_frame,
    data.frame(
      model = "Lasso favoring",
      modalities = "Clinical + Mutation",
      cancer = cancer,
      concordance = unlist(sapply(
        readRDS(here::here("data", "benchmarks", cancer, "glmnet_favor_10_stratified_clinical_mut.rds")),
        function(x) x$concordance
      ) %>% replace_na(
        0.5
      ))
    )
  )

  plt_frame <- rbind(
    plt_frame,
    data.frame(
      model = "Lasso favoring",
      modalities = "Clinical + Methylation",
      cancer = cancer,
      concordance = unlist(sapply(
        readRDS(here::here("data", "benchmarks", cancer, "glmnet_favor_10_stratified_clinical_meth.rds")),
        function(x) x$concordance
      ) %>% replace_na(
        0.5
      ))
    )
  )

  plt_frame <- rbind(
    plt_frame,
    data.frame(
      model = "Lasso favoring",
      modalities = "Clinical + RPPA",
      cancer = cancer,
      concordance = unlist(sapply(
        readRDS(here::here("data", "benchmarks", cancer, "glmnet_favor_10_stratified_clinical_rppa.rds")),
        function(x) x$concordance
      ) %>% replace_na(
        0.5
      ))
    )
  )

  plt_frame <- rbind(
    plt_frame,
    data.frame(
      model = "Lasso favoring",
      modalities = "Clinical + miRNA",
      cancer = cancer,
      concordance = unlist(sapply(
        readRDS(here::here("data", "benchmarks", cancer, "glmnet_favor_10_stratified_clinical_mirna.rds")),
        function(x) x$concordance
      ) %>% replace_na(
        0.5
      ))
    )
  )

  plt_frame <- rbind(
    plt_frame,
    data.frame(
      model = "Lasso favoring",
      modalities = "Clinical + CNV",
      cancer = cancer,
      concordance = unlist(sapply(
        readRDS(here::here("data", "benchmarks", cancer, "glmnet_favor_10_stratified_clinical_cnv.rds")),
        function(x) x$concordance
      ) %>% replace_na(
        0.5
      ))
    )
  )

  plt_frame <- rbind(
    plt_frame,
    data.frame(
      model = "Ridge",
      modalities = "Clinical",
      cancer = cancer,
      concordance = unlist(sapply(
        readRDS(here::here("data", "benchmarks", cancer, "clinical_ridge_10_stratified_all.rds")),
        function(x) x$concordance
      ) %>% replace_na(
        0.5
      ))
    )
  )
}

plt_frame %>%
  group_by(model, modalities) %>%
  summarise(mean = paste0(round(mean(concordance), 3), " (", round(sd(concordance), 3), ")")) %>%
  arrange(desc(mean)) %>%
  select(model, mean, modalities) %>%
  pivot_wider(
    names_from = modalities,
    values_from = mean
  ) %>%
  write_csv(
    here::here(
      "tables", "table_3.csv"
    )
  )
```
