## Supplementary material for "Clinically driven knowledge distillation for sparsifying high-dimensional multi-omics survival models": Code: reproduce_table_4.R

```
library(here)
library(rjson)
library(readr)
library(dplyr)
library(ggplot2)
library(tidyr)

config <- rjson::fromJSON(
  file = here::here("config", "config.json")
)
plt_frame <- data.frame()

for (cancer in config$cancers) {
  plt_frame <- rbind(
    plt_frame,
    data.frame(
      features = "KD favoring (BF)",
      model = "Ridge",
      cancer = cancer,
      concordance = unlist(sapply(
        readRDS(here::here("data", "benchmarks", cancer, "ridge_10_stratified_all_blockforest_feature_selected.rds")),
        function(x) x$concordance
      ))
    )
  )


  plt_frame <- rbind(
    plt_frame,
    data.frame(
      features = "KD favoring (RBF)",
      model = "Ridge",
      cancer = cancer,
      concordance = unlist(sapply(
        readRDS(here::here("data", "benchmarks", cancer, "ridge_10_stratified_all_rb_favoring_feature_selected.rds")),
        function(x) x$concordance
      ))
    )
  )


  plt_frame <- rbind(
    plt_frame,
    data.frame(
      features = "KD favoring (HSAE)",
      model = "Ridge",
      cancer = cancer,
      concordance = unlist(sapply(
        readRDS(here::here("data", "benchmarks", cancer, "ridge_10_stratified_all_hierarchical_sae_feature_selected.rds")),
        function(x) x$concordance
      ))
    )
  )


  plt_frame <- rbind(
    plt_frame,
    data.frame(
      features = "Lasso favoring",
      model = "Ridge",
      cancer = cancer,
      concordance = sapply(
        readRDS(here::here("data", "benchmarks", cancer, "ridge_10_stratified_all_glmnet_favoring_feature_selected.rds")),
        function(x) x$concordance
      ) %>% unlist() %>% replace_na(
        0.5
      )
    )
  )

  plt_frame <- rbind(
    plt_frame,
    data.frame(
      features = "prioritylasso",
      model = "Ridge",
      cancer = cancer,
      concordance = sapply(
        readRDS(here::here("data", "benchmarks", cancer, "ridge_10_stratified_all_prioritylasso_feature_selected.rds")),
        function(x) x$concordance
      ) %>% unlist() %>% replace_na(
        0.5
      )
    )
  )

  plt_frame <- rbind(
    plt_frame,
    data.frame(
      features = "Clinical-only",
      model = "Ridge",
      cancer = cancer,
      concordance = sapply(
        readRDS(here::here("data", "benchmarks", cancer, "clinical_ridge_10_stratified_all.rds")),
        function(x) x$concordance
      ) %>% unlist() %>% replace_na(
        0.5
      )
    )
  )
  plt_frame <- rbind(
    plt_frame,
    data.frame(
      features = "All",
      model = "prioritylasso",
      cancer = cancer,
      concordance = sapply(
        readRDS(here::here("data", "benchmarks", cancer, "prioritylasso_10_stratified_all.rds")),
        function(x) x$concordance
      )
    )
  )

  plt_frame <- rbind(
    plt_frame,
    data.frame(
      model = "Lasso favoring",
      features = "All",
      cancer = cancer,
      concordance = sapply(
        readRDS(here::here("data", "benchmarks", cancer, "glmnet_favor_10_stratified_all.rds")),
        function(x) x$concordance
      )
    )
  )

  plt_frame <- rbind(
    plt_frame,
    data.frame(
      model = "KD favoring (RBF)",
      cancer = cancer,
      features = "All",
      concordance = unlist(sapply(
        readRDS(here::here("data", "benchmarks", cancer, "randomblock_favoring_distilled_all.rds")),
        function(x) x[[4]]
      ) %>% replace_na(
        0.5
      ))
    )
  )

  plt_frame <- rbind(
    plt_frame,
    data.frame(
      model = "KD favoring (HSAE)",
      cancer = cancer,
      features = "All",
      concordance = unlist(sapply(
        readRDS(here::here("data", "benchmarks", cancer, "hierarchical_sae_distilled_all.rds")),
        function(x) x[[4]]
      ) %>% replace_na(
        0.5
      ))
    )
  )

  plt_frame <- rbind(
    plt_frame,
    data.frame(
      model = "KD favoring (BF)",
      features = "All",
      cancer = cancer,
      concordance = unlist(sapply(
        readRDS(here::here("data", "benchmarks", cancer, "blockforest_distilled_all.rds")),
        function(x) x[[4]]
      ) %>% replace_na(
        0.5
      ))
    )
  )
}


plt_frame %>%
  filter(features != "All") %>%
  group_by(model, features) %>%
  summarise(
    mean = mean(concordance),
    mean_conc = paste0(
      round(mean(concordance), 3),
      " (", round(sd(concordance), 3), ")"
    )
  ) %>%
  arrange(desc(mean)) -> tmp_1

plt_frame %>%
  filter(features == "All") %>%
  group_by(model, features) %>%
  summarise(
    mean = mean(concordance),
    mean_conc = paste0(
      round(mean(concordance), 3),
      " (", round(sd(concordance), 3), ")"
    )
  ) %>%
  arrange(desc(mean)) -> tmp_2

tmp_3 <- tmp_1[-grep("Clinical", tmp_1$features), ]
tmp_3$diff_clinical <- sapply(
  plt_frame %>% filter(features != "All" & features != "Clinical-only") %>% pull(features) %>% unique(),
  function(x) {
    paste0(
      round(mean(
        plt_frame %>% filter(features == x) %>% pull(concordance)
          - plt_frame %>% filter(features == "Clinical-only") %>% pull(concordance)
      ), 3),
      " (",
      round(sd(
        plt_frame %>% filter(features == x) %>% pull(concordance)
          - plt_frame %>% filter(features == "Clinical-only") %>% pull(concordance)
      ), 3), ")"
    )
  }
)
tmp_3$diff_scratch <- sapply(
  plt_frame %>% filter(features != "All" & features != "Clinical-only") %>% pull(features) %>% unique(),
  function(x) {
    paste0(
      round(mean(
        plt_frame %>% filter(features == x) %>% pull(concordance)
          - plt_frame %>% filter(model == x) %>% pull(concordance)
      ), 3),
      " (",
      round(sd(
        plt_frame %>% filter(features == x) %>% pull(concordance)
          - plt_frame %>% filter(model == x) %>% pull(concordance)
      ), 3), ")"
    )
  }
)

colnames(2:ncol(tmp_3)) <- c("Feature set from", "Mean concordance", "Delta clinical", "Delta originally trained model")
tmp[, -1] %>% write_csv(
  here::here(
    "tables", "table_4.csv"
  )
)
```
