## Supplementary material for "Clinically driven knowledge distillation for sparsifying high-dimensional multi-omics survival models": Code: utils.py

```
#!/usr/bin/env python
# -*- coding: utf-8 -*-

import os
import sys

module_path = os.path.abspath(os.path.join("./src/"))
if module_path not in sys.path:
    sys.path.append(module_path)

import random

import numpy as np
import torch
from sklearn.model_selection import StratifiedKFold, StratifiedShuffleSplit
from sklearn.model_selection._split import _RepeatedSplits
from skorch.callbacks import Callback
from skorch.dataset import CVSplit, get_len
from skorch.utils import to_numpy
from torch import nn

# All code adapted from:
# Wissel, D., Rowson, D., and Boeva, V. Hierarchical autoencoder-based integration improves performance in multi-omics cancer survival models through soft modality selection. bioRxiv, 2022.

# Adapted from https://github.com/pytorch/pytorch/issues/7068.
def seed_torch(seed=42):
    """Sets all seeds within torch and adjacent libraries.

    Args:
        seed: Random seed to be used by the seeding functions.

    Returns:
        None
    """
    random.seed(seed)
    os.environ["PYTHONHASHSEED"] = str(seed)
    np.random.seed(seed)
    torch.manual_seed(seed)
    torch.cuda.manual_seed(seed)
    torch.cuda.manual_seed_all(seed)
    torch.backends.cudnn.benchmark = False
    torch.backends.cudnn.deterministic = True
    return None


# Adapted from https://github.com/skorch-dev/skorch/issues/280
class FixRandomSeed(Callback):
    """Ensure reproducibility within skorch by setting all seeds.

    Attributes:
        seed: Random seed to be used by the seeding functions.

    """

    def __init__(self, seed=42):
        self.seed = seed

    def initialize(self):
        seed_torch(self.seed)


# Adapted from Tong, Li, et al. "Deep learning based feature-level integration of multi-omics data for breast cancer patients survival analysis." BMC medical informatics and decision making 20.1 (2020): 1-12.
# https://github.com/tongli1210/BreastCancerSurvivalIntegration/blob/master/src/models/loss_survival.py
def get_R_matrix(survival_time):
    """
    Create an indicator matrix of risk sets, where T_j >= T_i.

    Input:
        survival_time: a Pytorch tensor that the number of rows is equal top the number of samples
    Output:
        indicator matrix: an indicator matrix
    """
    batch_length = survival_time.shape[0]
    R_matrix = np.zeros([batch_length, batch_length], dtype=int)
    for i in range(batch_length):
        for j in range(batch_length):
            R_matrix[i, j] = survival_time[j] >= survival_time[i]
    return R_matrix


# Adapted from Tong, Li, et al. "Deep learning based feature-level integration of multi-omics data for breast cancer patients survival analysis." BMC medical informatics and decision making 20.1 (2020): 1-12.
# https://github.com/tongli1210/BreastCancerSurvivalIntegration/blob/master/src/models/loss_survival.py
def neg_par_log_likelihood(
    pred,
    survival_time,
    survival_event,
    sample_weight,
    cuda=False,
):
    """
    Calculate the average Cox negative partial log-likelihood
    Input:
        pred: linear predictors from trained model.
        survival_time: survival time from ground truth
        survival_event: survival event from ground truth: 1 for event
                and 0 for censored
    Output:
        cost: the survival cost to be minimized
    """
    sample_weight = torch.unsqueeze(sample_weight, 1)
    survival_event = torch.tensor(survival_event)
    survival_time = torch.tensor(survival_time)
    n_observed = survival_event.sum(0)
    if not n_observed:
        # Return zero loss if there are no events
        # within a batch.
        return torch.tensor(0.0)
    R_matrix = get_R_matrix(survival_time)
    R_matrix = torch.Tensor(R_matrix)
    if cuda:
        R_matrix = R_matrix.cuda()
    risk_set_sum = R_matrix.mm(torch.exp(pred))
    diff = pred - torch.log(risk_set_sum)
    survival_event = torch.reshape(
        survival_event, (survival_event.shape[0], 1)
    )
    sum_diff_in_observed = (
        torch.transpose(diff * sample_weight, 0, 1).float().mm(survival_event)
    )
    loss = (-(sum_diff_in_observed) / n_observed).reshape((-1,))
    return loss


class hierarchical_sae_criterion(nn.Module):
    """torch criterion to calculate various losses needed by the
    HierarchicalSAE class.

    Attributes:
        None
    """

    def forward(self, input, target, sample_weight, n_blocks):
        """Calculate losses needed for backprop of the HierachicalSAE
        class.

        Args:
            input: Predicted partial hazard vector (nx1).
            target: Survival target vector of the format 'event|time' (nx1).
            sample_weight: Sample weight vector (nx1).
            n_blocks: Number of modalities inputted to HierarchicalSAE.


        """
        mse = nn.MSELoss(reduction="none")
        losses = {
            # Doing this matrix multiplication implicitly
            # multiplies the reconstruction loss
            # by the dataset size, i.e., the batch size.
            "common_reconstruction": torch.mean(
                torch.unsqueeze(sample_weight, dim=0)
                .float()
                .mm(mse(input[2], input[1]))
            ),
            "cox": neg_par_log_likelihood(
                input[0],
                np.array([str.rsplit(i, "|")[1] for i in target]).astype(
                    np.float32
                ),
                np.array([str.rsplit(i, "|")[0] for i in target]).astype(
                    np.float32
                ),
                sample_weight,
            ),
        }
        for ix in range(n_blocks):
            # Doing this matrix multiplication implicitly
            # multiplies the reconstruction loss
            # by the dataset size, i.e., the batch size.
            losses[f"block_{ix}_reconstruction"] = torch.mean(
                torch.unsqueeze(sample_weight, dim=0)
                .float()
                .mm(mse(input[4][ix], input[3][ix]))
            )
            losses[f"block_{ix}_hazard"] = neg_par_log_likelihood(
                input[-2][ix],
                np.array([str.rsplit(i, "|")[1] for i in target]).astype(
                    np.float32
                ),
                np.array([str.rsplit(i, "|")[0] for i in target]).astype(
                    np.float32
                ),
                sample_weight,
            )
        return losses


class StratifiedSurvivalKFold(StratifiedKFold):
    """Adapt `StratifiedKFold` to make it usable with our adapted
    survival target string format.

    For further documentation, please refer to the `StratifiedKFold`
    documentation, as the only changes made were to adapt the string
    target format.
    """

    def _make_test_folds(self, X, y=None):
        if y is not None and isinstance(y, np.ndarray):
            # Handle string target by selecting out only the event
            # to stratify on.
            if not y.dtype == np.dtype("float32"):
                y = np.array([str.rsplit(i, "|")[0] for i in y]).astype(
                    np.float32
                )
        return super()._make_test_folds(X=X, y=y)

    def _iter_test_masks(self, X, y=None, groups=None):
        if y is not None and isinstance(y, np.ndarray):
            # Handle string target by selecting out only the event
            # to stratify on.
            if not y.dtype == np.dtype("float32"):
                y = np.array([str.rsplit(i, "|")[0] for i in y]).astype(
                    np.float32
                )
        return super()._iter_test_masks(X, y=y)

    def split(self, X, y, groups=None):
        return super().split(X=X, y=y, groups=groups)


class StratifiedSurvivalShuffleSplit(StratifiedShuffleSplit):
    """Adapt `StratifiedShuffleSplit` to make it usable with our adapted
    survival target string format.

    For further documentation, please refer to the `StratifiedShuffleSplit`
    documentation, as the only changes made were to adapt the string
    target format.
    """

    def split(self, X, y, groups=None):
        # Handle string target by selecting out only the event
        # to stratify on.
        y = np.array([str.rsplit(i, "|")[0] for i in y]).astype(np.float32)
        return super().split(X, y, groups)


class RepeatedStratifiedSurvivalKFold(_RepeatedSplits):
    """Adapt `_RepeatedSplits` to make it usable with our adapted
    survival target string format.

    For further documentation, please refer to the `_RepeatedSplits`
    documentation, as the only changes made were to adapt the string
    target format.
    """

    def __init__(self, *, n_splits=5, n_repeats=10, random_state=None):
        # Handle string target format by using previously defined
        # `StratifiedSurvivalKFold` class, which handles survival
        # string format.
        super().__init__(
            StratifiedSurvivalKFold,
            n_repeats=n_repeats,
            random_state=random_state,
            n_splits=n_splits,
        )


class StratifiedSkorchSurvivalSplit(CVSplit):
    """Adapt `CVSplit` to make it usable with our adapted
    survival target string format.

    For further documentation, please refer to the `CVSplit`
    documentation, as the only changes made were to adapt the string
    target format.
    """

    def __call__(self, dataset, y=None, groups=None):
        if y is not None:
            # Handle string target by selecting out only the event
            # to stratify on.
            y = np.array([str.rsplit(i, "|")[0] for i in y]).astype(np.float32)

        bad_y_error = ValueError(
            "Stratified CV requires explicitly passing a suitable y."
        )

        if (y is None) and self.stratified:
            raise bad_y_error

        cv = self.check_cv(y)
        if self.stratified and not self._is_stratified(cv):
            raise bad_y_error

        # pylint: disable=invalid-name
        len_dataset = get_len(dataset)
        if y is not None:
            len_y = get_len(y)
            if len_dataset != len_y:
                raise ValueError(
                    "Cannot perform a CV split if dataset and y "
                    "have different lengths."
                )

        args = (np.arange(len_dataset),)
        if self._is_stratified(cv):
            args = args + (to_numpy(y),)

        idx_train, idx_valid = next(iter(cv.split(*args, groups=groups)))
        dataset_train = torch.utils.data.Subset(dataset, idx_train)
        dataset_valid = torch.utils.data.Subset(dataset, idx_valid)
        return dataset_train, dataset_valid
```
