## Supplementary material for "Clinically driven knowledge distillation for sparsifying high-dimensional multi-omics survival models": Code: utils.R

```
library(here)
library(stringr)

format_splits <- function(raw_splits) {
  if (any(is.na(raw_splits))) {
    apply(data.frame(raw_splits), 1, function(x) unname(x[!is.na(x)]) + 1)
  } else {
    x <- unname(as.matrix(raw_splits)) + 1
    split(x, row(x))
  }
}

format_features <- function(raw_feature_list) {
  unname(
    sapply(
      strsplit(
        raw_feature_list, "',|\","
      )[[1]], function(x) {
        str_replace_all(str_replace_all(str_replace_all(str_replace_all(str_replace_all(
          str_replace_all(
            str_replace_all(str_replace_all(str_replace_all(x, "\\['", ""), "^ '", ""), '^ \\"', ""), '\\"', ""
          ),
          "^\'", ""
        ), "\'$", ""), "'\\]", ""), "\\[", "\\\\["), "\\]", "\\\\]")
      }
    )
  )
}
```
